# Excessive Dietary Fructose Aggravates Heart Failure via Impairing Myocardial Fatty Acid Oxidation Metabolism in Diet Induced Obese Mouse

**DOI:** 10.1101/2024.05.22.595423

**Authors:** Yufeng Zhang, Yuzhou Xue, Qifan Gong, Jingshen Xu, Shuaikang Wang, Min Zhu, Jinhui Wang, Zhiqiang Song, Shuxian Zhang, Hu Wang, Ling Jin, Kun Hua, Xiubin Yang, Jianping Li, Jin Li, Ming Xu, He Huang

## Abstract

**Background:** An unhealthy diet and a sedentary lifestyle increase the prevalence of cardiometabolic syndrome. Several cardiovascular diseases (CVDs) have been highly linked to excessive added sugar intake, which alters whole-body metabolism, including heart tissue. However, whether specific added sugars can cause and aggravate severe heart dysfunction is still unknown.

**Methods:** We examined the association between CVDs and added sugar intake using statistical analyses and Mendelian randomization (MR). Then, we evaluated the effect of added sugar on mouse heart function employing a diet-induced obese (DIO) model with transverse aortic constriction (TAC) challenge. We measured the fatty acid and fructose metabolic flux in mouse hearts employing a mass spectrometry-based *in vivo* double stable isotopic labeling (DSIL) technique. The results of animal models were also confirmed in aortic stenosis (AS) patient samples.

**Results:** Statistical analyses and MR estimation on public databases indicated that added sugars, especially fructose, are associated with a high risk of heart failure. Feeding on either ingested or drunk fructose could aggravate heart failure and diastolic and systolic dysfunction in TAC challenged DIO mice. Mechanistically, excessive dietary intake of fructose could suppress heart fatty acid oxidation (FAO) metabolism via both shrinkage of the flux rate of FAO and inhibition of the AMPK-ACC axis. Activation of AMPK or deactivation of ACC could limit such heart dysfunction and myocardial hypertrophy. We also obtained plasma from 27 patients with AS and determined that the high fructose level instead of glucose or sucrose was linked to left ventricular ejection fractions (LVEF) and fractional shortening (FS) decline.

**Conclusions:** Findings from epidemiological statistical analyses and investigations of animal models suggested the harmful effect of excessive fructose intake on heart function. Fructose could directly alter heart metabolism by suppressing FAO. Our results implied that targeting AMPK-ACC mediation could effectively attenuate excessive fructose-induced heart failure in DIO mice.

## Introduction

Cardiometabolic syndrome (CMS) has become a primary epidemiological burden in modern industrialized countries in the last decade^1^. Evidence has suggested that unhealthy diet patterns and a sedentary lifestyle are two predominant causes of CMS and elevate the risk of coronary heart disease, ischemic stroke, and cardiovascular mortality^2–5^. Among unhealthy diet patterns, excessive added sugar, including high fructose corn syrup-added beverages, has been identified as a cause of the high prevalence of multiple cardiovascular diseases (CVDs)^5–7^. However, many studies focused on added sugar-induced hepatic lipogenesis and inflammation, in which heart dysfunction has been considered a complication^6^. The direct influence of added sugar on heart function and myocardial metabolism was overlooked. Whether such excessive added sugar intake in an unhealthy diet can directly produce CVD is still unclear.

Cardiac metabolism alterations could cause chronic pathophysiological circumstances^8,9^. Typically, a healthy heart favors free fatty acid consumption as the primary energy substrate instead of glucose. Approximately 40% to 60% of cardiac energy is produced from fatty acid oxidization, which consumes the most oxygen delivered to the heart^10^. The changes in cardiac fatty acid and carbohydrate metabolism could contribute to the progression of cardiac hypertrophy, hypertension, and recurrent ischemic insult^11–13^. Additionally, using fatty acids as fuels in cardiomyocytes is highly correlated to heart health. For instance, the rate of myocardial fatty acid metabolism declines as the severity of heart failure progresses (ejection fraction below 50%)^14,15^.

In this study, we aimed to assess the influence of added sugar on cardiac metabolism and heart failure progression. Combing our results of clinical analyses and animal models, we discovered that added sugar, especially excessive fructose intake, could produce heart failure by rapidly altering fatty acid metabolism in diet-induced obese (DIO) mice. The stable isotope labeling-based metabolic flux analysis revealed competitive catabolism of fructose in mouse hearts could limit fatty acid oxidation (FAO), and excessive fructose could also mediate FAO via the AMPK-ACC axis. Our study suggested that the excessive dietary intake of fructose is a risk factor for heart failure. Control of dietary fructose intake may be an important strategy to prevent heart failure progression.

## Methods

The *Supplementary Methods* offers an expanded methods section. The data supporting this study’s findings are available from the corresponding author upon reasonable request.

### Public Database Statistic Analyses

In this study, a meta-analysis was conducted on the National Health and Nutrition Examination Survey (NHANES) data (n = 91,351) from 2001 to 2018, representing nine cycles^16^. Based on exclusion criteria, a total of 40,058 participants were enrolled in our analysis (Figure S1). Nutrient intake from food was estimated based on the US Department of Agriculture Food and Nutrient Database for Dietary Studies (FNDDS)^17^. The macronutrient (carbohydrate, fats, and protein) energy (%E) intake was characterized by separating energy from nutrients by the total daily energy intake (for instance, carbohydrate %E = ([carbohydrate (g) × 4]/total energy intake [kcal]) × 100). Hypertension was identified according to three aspects: (1) have diagnosed with hypertension by a doctor, (2) took anti-hypertensive drugs, or (3) had at least three blood pressure measurements of: (systolic blood pressure ≥ 140 mmHg or diastolic blood pressure ≥ 90 mmHg). Coronary heart disease and congestive heart failure were identified from a disease questionnaire. The covariates encompassed age, gender, race, educational level (< 9, 9-12, or > 12 years), marital status, body mass index, smoking status, diabetic status, antihypertensive medication use, Healthy Eating Index (HEI) score, total serum cholesterol, serum glucose, and total energy intake. Across all analyses, two-year MEC exam weights were employed for all participant estimations, accounting for sampling design complexity.

### Two-sample Mendelian Randomization Analysis

We assessed the causal relationship between blood sugars (sucrose, glucose, and fructose) and the risk of cardiovascular diseases using a two-sample Mendelian randomization (MR) estimation approach (Figure S2). Genetic information for all blood metabolites except sucrose was obtained from the Shin *et al.* study, in which a total of 7,824 adults of European ancestry were included, and approximately 2.1 million SNPs were identified^18^. While the genetic information on blood sucrose was accessed from the study of Long *et al.*^19^. This study analyzed serum samples from 1960 subjects of European descent and approximately 11.3 million variants. The GWAS statistics for cardiovascular diseases (heart failure, hypertension, and coronary artery disease) were harvested from the EBI database (publicly available at https://gwas.mrcieu.ac.uk/). Detailed information about diagnosis criteria, demography, and quality control can be obtained from original studies^20–23^.

Instrument identification was performed via a series of steps to select eligible genetic variants. Firstly, we selected a *p*-value threshold less than 5×10^-5^ for the instrument SNP screen as previously described^24^. Secondly, linkage disequilibrium (LD) r^2^ < 0.01 within a 100 kb distance was conducted to identify independent SNPs. Additionally, F statistics were determined for each SNP to exclude weak instrument, and SNPs with F < 10 were discarded from downstream analysis. Finally, the extracted SNPs with strong association with outcomes (*p* < 0.005) were discarded, while the proxy SNPs for outcomes were interrogated with a minimum LD r^2^ > 0.8.

The primary analysis was characterized as a random-effect inverse variance weighted (IVW) method to identify causal relationships (*p* < 0.05). Furthermore, other MR methods, notably MR-PRESSO, Maximum likelihood-based, MR-Egger, and Weighted median, were complementary. Horizontal pleiotropy was examined via Egger intercept. The Cochran-Q test was conducted to evaluate the heterogeneity (*p* < 0.05 and I^2^ > 25% was recognized as heterogeneity). Leave-one-out (LOO) analysis was utilized to assess the influence of each exposure-associated variant on the IVW estimates. We conducted bidirectional MR to explore the causal effect of heart failure on blood sugars (mirroring parameters as above). All analyses were performed using the “TwosampleMR” package (Version 0.4.26) and the “MRPRESSO” package (Version 1.0) based on the R Program (Version 4.2.1).

### Patient Recruitment, Sample Collection, and Clinical Feature Measurement

Consecutive aortic valve stenosis patients (age > 18 years) who were diagnosed at Anzhen Hospital (from 2021.12 to 2022.03) were recruited for this study (Table S1). Patients with other cardiovascular diseases (including coronary heart disease, hyperlipidemia, and diabetes mellitus), kidney injury (estimated glomerular filtration rate, eGFR < 60 mL/min/1.73 m^2^), and digestive disease (gastritis, peptic ulcer, gastric cancer, pancreatic cancer, and other gastrointestinal tumors) were excluded from the analysis. Fasting blood was harvested in the first 24 hours after administration and stored at −80 ℃ before analyses. Transthoracic echocardiography was applied to evaluate the baseline cardiac function. The demographics, clinical data, and laboratory tests were prospectively collected in databases. The plasma fructose level was quantified by a fructose assay kit (ab83380, Abcam). The study received approval from the local institutional ethics committee (2021140X).

## Animal Studies

### Animal Feeding and Treatment

Six-week-old male C57BL/6J mice were obtained from GemPharmatech (Nanjing, China). Animals were housed under controlled temperature (23 °C) and light, with access to water and food *ad libitum*. The diet-induced obese (DIO) mouse was fed either a high-fat diet (Research Diets, D12492) or a high-fat, high-carbohydrate diet (SYSE BIO, TD120528). Thirty percent fructose drinking water for DIO mouse feeding was prepared. For chemical drug treatments, mice were administered daily intraperitoneal injections of 5-Aminoimidazole-4-carboxamide1-β-D-ribofuranoside (AICAR) (MedChemExpress, HY-13417) at a dose of 500 mg/kg, ACC inhibitor (CP-640186) (MedChemExpress, HY-15259) at a dose of 25 mg/kg or saline as a control from one-week post-TAC surgery. All mice were fasted for six hours before dissection.

### Transverse Aortic Constriction

Surgery was conducted as previously described^25^. Specifically, mice were anesthetized using xylazine (5 mg/kg) via intraperitoneal injection. Thereafter, a midline sternal incision was made to expose the thymus and arteries. Following this, the thymus was gently separated from the aortic arch, and transverse aortic constriction (TAC) was performed by performing a ligature with a 7-0 nylon suture against a 27 G needle. The skin was closed with a 3-0 suture. The sham group was subjected to the same procedure without aorta ligation.

### Echocardiogram Evaluation

Echocardiographic analysis was conducted using a Vevo 2100 High-Resolution Digital-Imaging System (Visual Sonics, Toronto, Canada) equipped with an MS400 transducer. Mice were anesthetized employing isoflurane in 100% oxygen (4% for induction and 1.5% for maintenance) and secured with surgical tape in a supine position. Heart rates were maintained at 350-450 beats per minute for diastolic function, while mice were conscious for systolic function. The cardiac function was evaluated at baseline, 2, 4, and 6 weeks following TAC.

### Stable Isotopic Metabolic Labeling

On the day of injection, mice were transferred to new cages without food around 9 AM (beginning of their sleep cycle), and the injection was conducted around 3 PM. The oleic acid solution was prepared by combining oleic acid with bovine serum albumin (fatty acid-free) (Beyotime, ST025) at a molar ratio of 4:1 and dissolved in sterile physiological saline. Two injection solutions were prepared for double stable isotopic labeling (DSIL) experiment: oleic acid-d_2_ (OA-d_2_) + ^12^C6-fructose and ^13^C6-fructose + oleic acid (OA), by dissolving 200 mM ^12^C6-fructose (Solarbio, F8100) or ^13^C6-fructose (CIL, CLM-1553) in 15 mM oleic acid-d2 (MedChemExpress, HY-N1446S1) solution or unlabeled oleic acid (MCE HY-N1446) solution, respectively.

To account for natural abundance isotope interference, a control group lacking injection served as time zero. These mice were sacrificed, and their heart tissues were harvested for further analysis. To examine the metabolic flux of free fatty acids or fructose in the hearts of mice, each mouse with a different diet received a tail vein injection of OA-d2 + ^12^C6-fructose solution or ^13^C6-fructose + OA. The injection was administered at a rate of 3 μL/g of body weight. The mice were then sacrificed at 0 minutes, 5 minutes, 10 minutes, 20 minutes, and 30 minutes, and heart tissues were obtained for subsequent analysis.

### Metabolic Flux Analysis

Mouse heart samples were ground using liquid nitrogen immediately following removal. A sample of 1.5 mL of extractant (60 mM trichloroacetic acid at a 50:50 ratio with water: methanol, *v:v*) was mixed with the ground heart. This mixture was centrifuged at 4 °C at 15,000 × g for 10 minutes, and the supernatant was collected in a 2 mL centrifuge tube. Solid-phase extraction (SPE) was performed using a vacuum extraction apparatus. Generally, the extraction column (Waters, WAT094225) was activated using 1 mL of methanol and 1 mL of water. The collected supernatant was then added to the extraction column. After rinsing the column with 1 mL of water to remove impurities, the analytes were eluted three times alongside 0.5 mL of methanol solution containing 20 mM ammonium acetate. The eluate (1.5 mL in total) was pooled and dried under a stream of nitrogen, followed by immediate analysis by mass spectrometry (MS). Detailed methods for MS analysis and metabolic flux rate calculations are outlined in the *Supplementary Methods*.

### Immunoblotting

To evaluate the levels of p-AMPKα, p-ACC, CPT1A, and CPT1B in the heart, freshly isolated heart tissues were mixed with ice-cold RIPA lysis buffer (supplemented with protease inhibitor cocktail, 20 μL/mg heart weight). The lysates were homogenized and centrifuged at 20,000 × g for 10 minutes at 4 °C, and a fourfold volume of 5 × SDS-PAGE sample loading buffer was added to the supernatant. The levels of proteins were then examined via immunoblotting (IB). The band intensities were quantified using Image J software (National Institutes of Health Freeware). Antibodies employed in immunoblotting are outlined in the *Supplementary Methods*.

### Statistical Analyses

All data produced in animal studies were evaluated with Mann-Whitney or Student’s *t-*test (GraphPad Prism 7), where noted. The significance level was set at \**p* < 0.05, \*\**p* < 0.01, or \*\*\**p* < 0.001 for all cases.

## Results

### Added dietary sugar is a risk factor for heart failure

To evaluate dietary behavior and CMS correlation, we analyzed the relationship of three macronutrients (carbohydrate, protein, and fat) with three primary CVD outcomes (heart failure, hypertension, and coronary heart disease) using the NHANES database. From the database, 1,260 (3.15%), 1,701 (4.25%), and 17,226 (4.25%) individuals were diagnosed with heart failure, coronary heart disease, and hypertension, respectively. As anticipated, participants suffering from obesity and overweight conditions were more likely to be associated with the three major cardiovascular diseases (Table S2). Once the total energy intake, body weight, and BMI were adjusted using multinomial logistics regression models, the additional sugar intake (teaspoon) was significantly associated with a high risk of heart failure (OR 1.002; 95%CI 1.000 to 1.003; *p* < 0.01; and OR 1.002; 95%CI 1.000 to 1.004; *p* < 0.05; Table 1). Similarly, the carbohydrate intake was more positively associated with heart failure (OR 1.008; 95%CI 1.001 to 1.016; *p* = 0.035) and coronary heart disease (OR 1.009; 95%CI 1.001 to 1.016; *p* = 0.027) than the protein or fat intake (Table S3). Total sugar intake was positively associated with heart failure (OR 1.002; 95%CI 1.000 to 1.003; *p* < 0.05) with adjusted BMI and energy intake (Table S4). These findings suggested that although obesity was highly associated with CVD risk, excessive added sugar intake could aggravate such risk, especially heart failure risk.

**Table 1:**
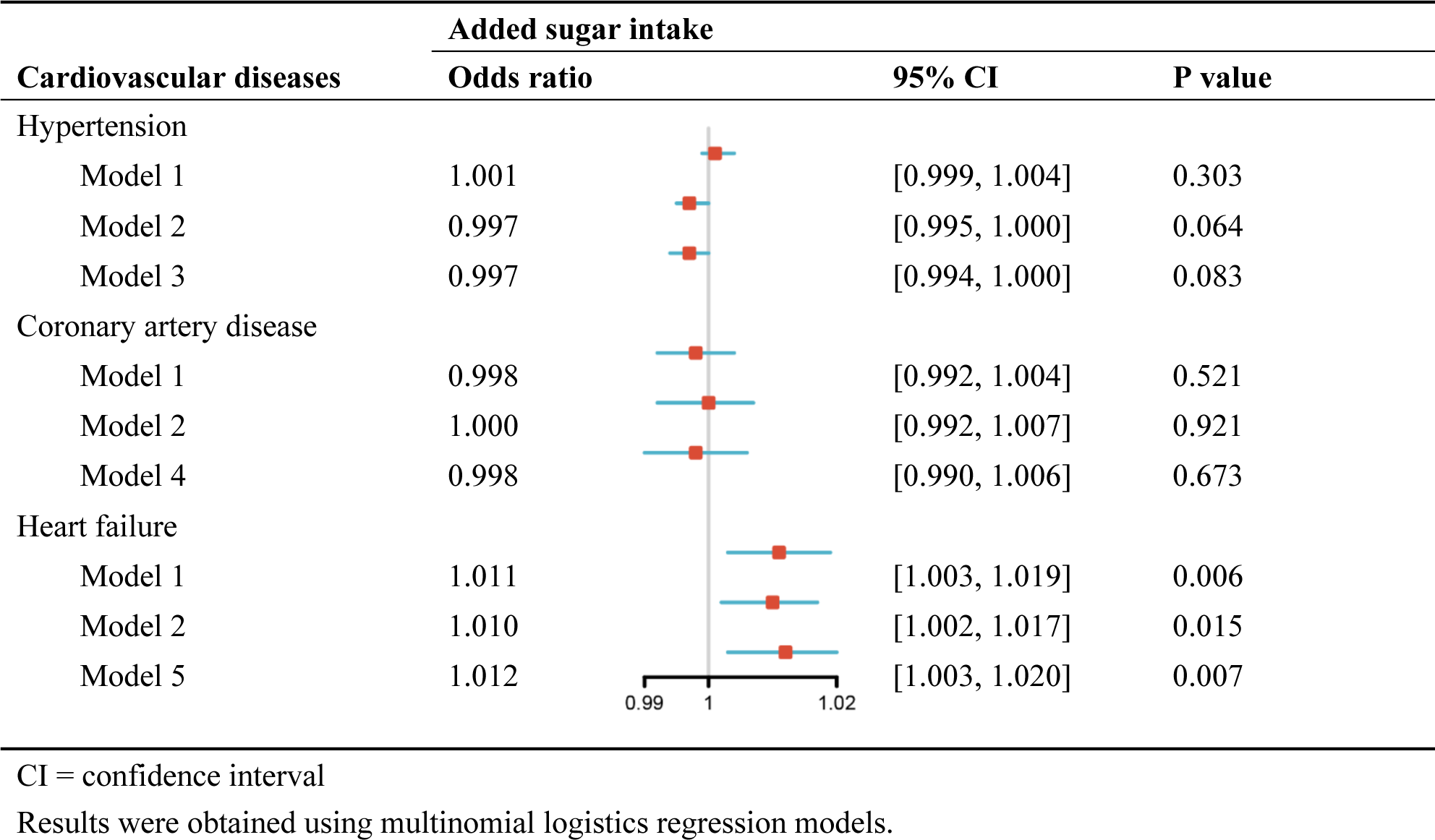
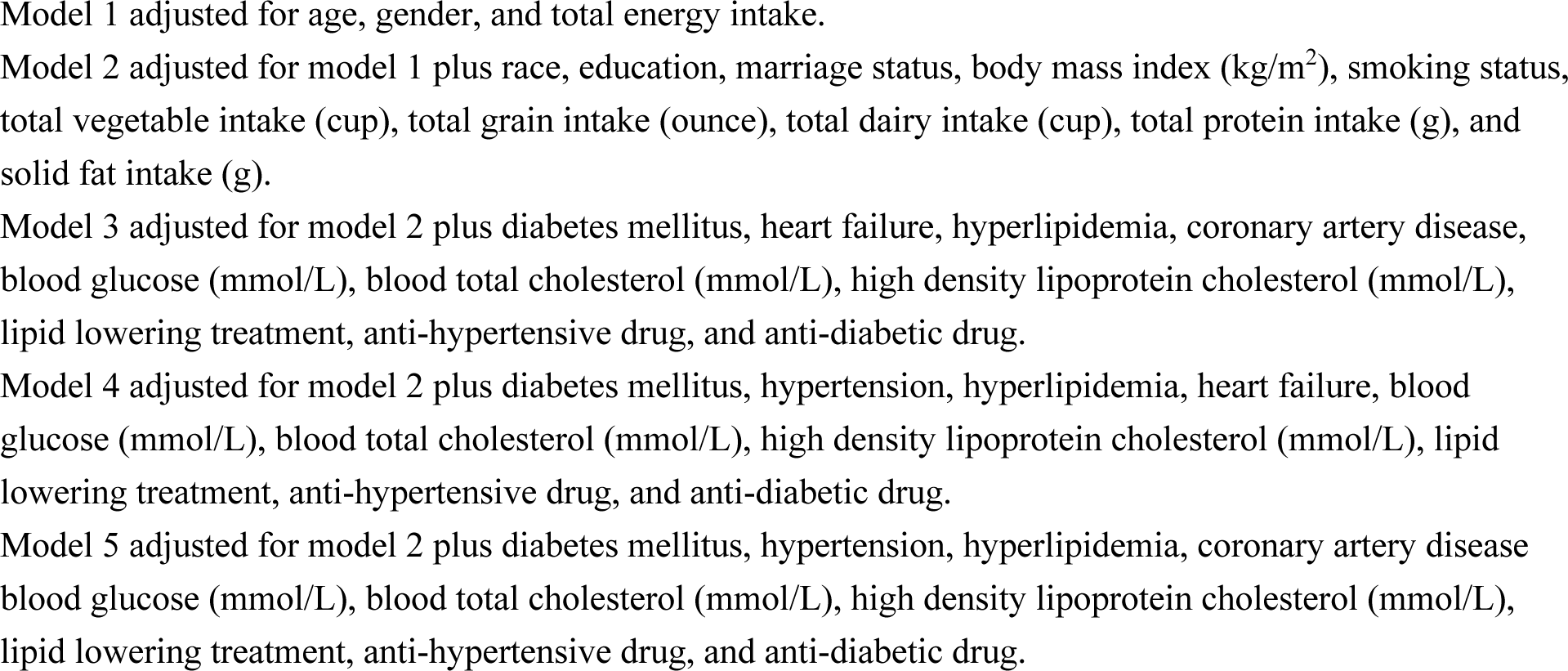
Relationship between added sugar intake (teaspoons) per day with various cardiovascular.

| Cardiovascular diseases | Added sugar intake |  |  | P value |
| --- | --- | --- | --- | --- |
|  | Odds ratio | 95% CI |  |  |
| Hypertension |  |  |  |  |
| Model 1 | 1.001 | [0.999, 1.004] |  | 0.303 |
| Model 2 | 0.997 | [0.995, 1.000] |  | 0.064 |
| Model 3 | 0.997 | [0.994, 1.000] |  | 0.083 |
| Coronary artery disease |  |  |  |  |
| Model 1 | 0.998 | [0.992, 1.004] |  | 0.521 |
| Model 2 | 1.000 | [0.992, 1.007] |  | 0.921 |
| Model 4 | 0.998 | [0.990, 1.006] |  | 0.673 |
| Heart failure |  |  |  |  |
| Model 1 | 1.011 | [1.003, 1.019] |  | 0.006 |
| Model 2 | 1.010 | [1.002, 1.017] |  | 0.015 |
| Model 5 | 1.012 | [1.003, 1.020] |  | 0.007 |

CI = confidence interval
Results were obtained using multinomial logistics regression models.
Model 1 adjusted for age, gender, and total energy intake.
Model 2 adjusted for model 1 plus race, education, marriage status, body mass index ( $\text{kg/m}^2$ ), smoking status, total vegetable intake (cup), total grain intake (ounce), total dairy intake (cup), total protein intake (g), and solid fat intake (g).
Model 3 adjusted for model 2 plus diabetes mellitus, heart failure, hyperlipidemia, coronary artery disease, blood glucose (mmol/L), blood total cholesterol (mmol/L), high density lipoprotein cholesterol (mmol/L), lipid lowering treatment, anti-hypertensive drug, and anti-diabetic drug.
Model 4 adjusted for model 2 plus diabetes mellitus, hypertension, hyperlipidemia, heart failure, blood glucose (mmol/L), blood total cholesterol (mmol/L), high density lipoprotein cholesterol (mmol/L), lipid lowering treatment, anti-hypertensive drug, and anti-diabetic drug.
Model 5 adjusted for model 2 plus diabetes mellitus, hypertension, hyperlipidemia, coronary artery disease, blood glucose (mmol/L), blood total cholesterol (mmol/L), high density lipoprotein cholesterol (mmol/L), lipid lowering treatment, anti-hypertensive drug, and anti-diabetic drug.

### Potential causal effects of blood fructose elevation on heart failure

Although we found a positive correlation between a high carbohydrate diet and heart failure, it was still necessary to comprehend which added sugar could produce CVD, specifically. Given the dietary consumption of added sugar typically encompassed sucrose, fructose, and glucose, we analyzed the statistical causality of these sugars on three CVDs (Figure S2). Inverse-variance weighted (IVW) identified that only a high blood fructose level induced heart failure (OR 1.216; 95%CI 1.041 to 1.420; *p* < 0.05; Figure 1A), but blood glucose (OR 1.008; 95%CI 0.803 to 1.265; *p* = 0.116) or sucrose (OR 0.996; 95%CI 0.991 to 1.001; *p* = 0.097) would not (Figure 1B and 1C). By performing sensitivity analyses, blood fructose was found to be involved in heart failure instead of glucose and sucrose. Four other MR models, including MR-PRESSO, Maximum likelihood-based, MR-Egger, and Weight Median tests, provided consistent direction and magnitude, which validated the robustness of causal association (Table S5-S7). For reverse MR analysis, genetic predisposition to heart failure was not linked to higher fructose (OR 1.00; 95%CI 0.98 to 1.02; *p* = 0.86) (Table S8). Furthermore, intercept analysis from MR-Egger indicated no risk of directional pleiotropy. Additionally, Cochran’s Q and I^2^ statistics indicated the absence of heterogeneity in the association between blood fructose and heart failure (Table S9). These findings implied that fructose, rather than other added sugars, would induce a higher risk of heart failure.

**Figure 1.**
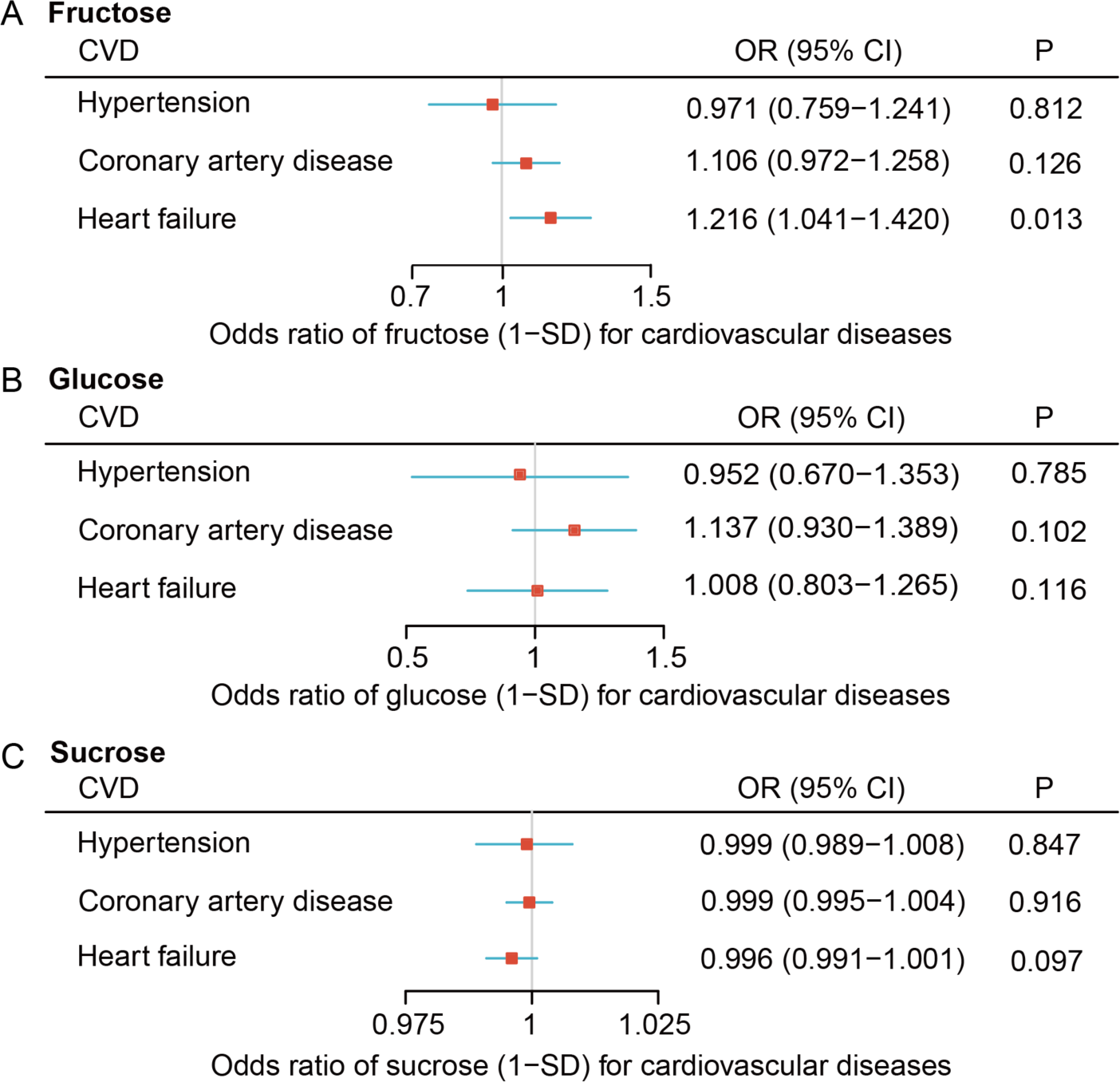
Mendelian randomization (MR) analyses of three commonly added sugars and three cardiovascular diseases indicate that (A) fructose had a higher causal effect on heart failure, but (B) glucose or (C) sucrose did not. MR estimates, and *p*-values were calculated using an inverse variance-weighted method with a random effects model. More details can be found in the *Methods* section.

### Excessive dietary fructose intake impairs cardiac function and aggravates heart morphological lesions in DIO mice

When high levels of fructose as an added sugar were established as a possible factor inducing heart failure epidemiologically, we designed three different diets to treat mice and see if excessive dietary fructose could induce heart dysfunction in animal models. To mirror modern unhealthy diet patterns, we chose two types of fructose intake approaches, adding to metabolic research on diet-induced obese (DIO) mouse models^26^: high-fat diet treatment (regular DIO model), high-fat, high-carbohydrate diet (DIO model with excessive ingestion fructose; DIO+Carb), and HFD with additional fructose drinking water (DIO model with excessive drinking fructose; DIO+Fruc) treatments. Compared to normal HFD (60% fats and 20% carbohydrate in total calories), the DIO+Carb and DIO+Fruc diets offered more ingestion (42% fats and 57% carbohydrates) or drinking fructose, respectively. Such a relatively brief dietary treatment would not induce any inflammation response in mice (Figure S3). Feeding the mice with three different diets, we challenged them with transverse aortic constriction (TAC) after two weeks of dietary treatments (Figure 2A). As anticipated, mice suffered multiple cardiological dysfunctions following TAC within six weeks. The survival probability was decreased in DIO+Carb and DIO+Fruc groups (Figure S4A). Echocardiographic results also indicated the diastolic (E/E’) and systolic function (LVEF, FS) of DIO+Carb and DIO+Fruc mice were profoundly impaired compared to the regular DIO mice beginning two weeks after TAC (Figure 2B-D and Figure S4B-D). The 2D-echocardiography also indicated an increased LVPW,d in DIO+Fruc treated mice (*p* = 0.0081), but this LVPW,d difference was shrunk after the TAC challenge (Figure S4Ei-Eii). These findings indicated that either excessive eating or drinking of fructose could impair heart function in DIO mice.

**Figure 2.**
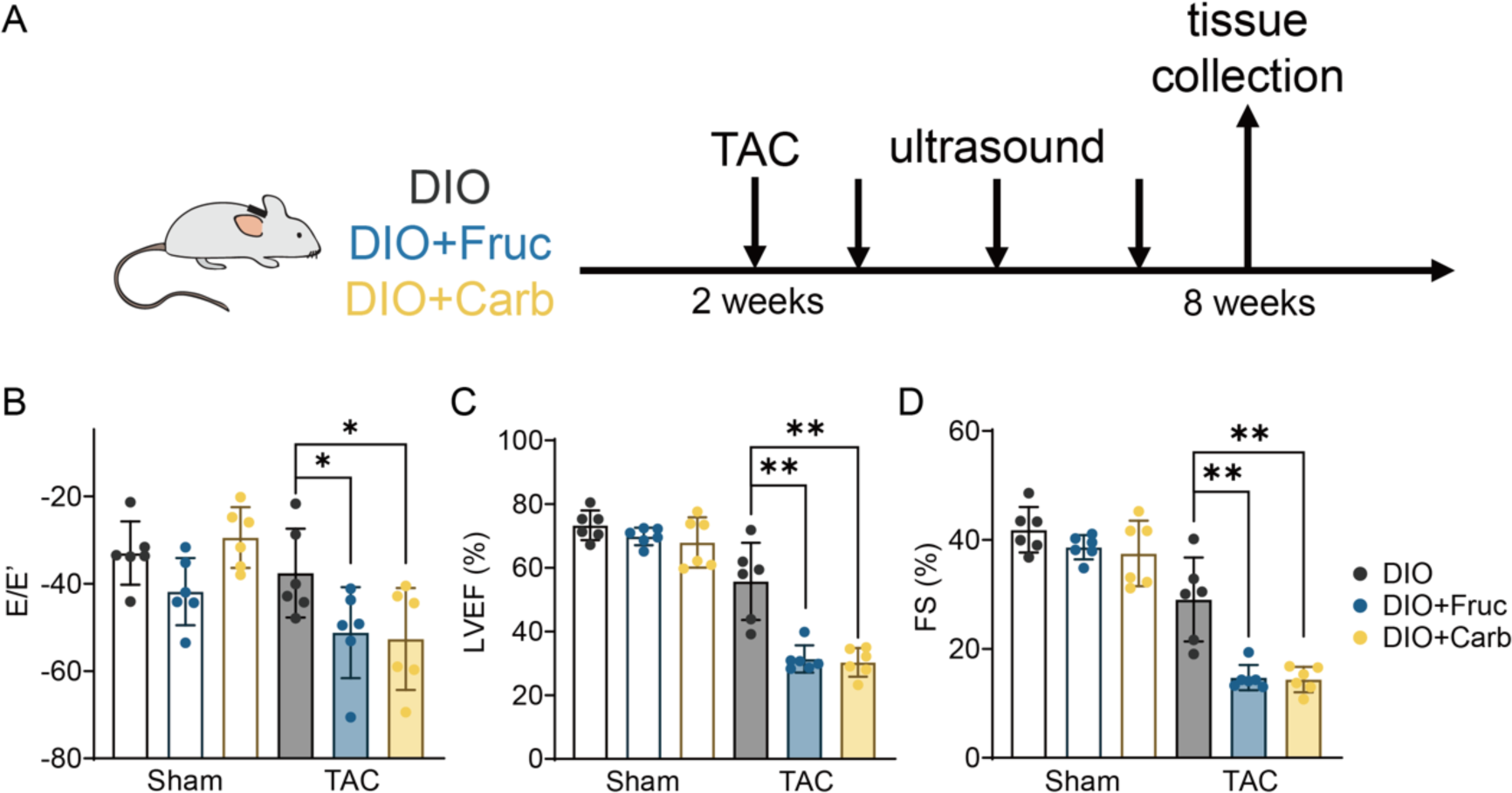
Excessive fructose impairs heart function in DIO mice with transverse aortic constriction (TAC) challenge. (A) Workflow of DIO mice group treatment. (B) Left ventricular ejection fractions (LVEF), (C) fractional shortening (FS), and (D) E/E’ were reduced following TAC surgery in DIO mice fed additional sugar (DIO+Carb) or drinking fructose (DIO+Fruc) (*vs.* DIO, \**p* < 0.05, \*\**p* < 0.01, n=6).

After monitoring the echocardiography of three diet-induced mice, they were sacrificed, and their heart weights were measured. As expected, the heart weight-to-tibia length ratio of both DIO+Fruc and DIO+Carb mice was significantly elevated following the TAC challenge (Figure 3A). Additionally, the histological findings of mice hearts indicated enlargement of both total heart tissue (Figure 3Bi, H&E staining) and cross-sectional cardiomyocyte area (Figure 3Bi-Bii, WGA staining) upon excessive fructose diet treatments. The PSR staining of heart tissue demonstrated that DIO+Fruc and DIO+Carb mice acquired larger interstitial and perivascular fibrosis than the normal DIO mice (Figure 3Ci-Cii). These findings indicated that excessive fructose could accelerate myocardial hypertrophy in DIO mice after the TAC challenge. When we observed the cardiomyocyte structure under electron microscopy, we were intrigued to find the accumulation number (Figure 3Di-Dii) and enlarged volume (Figure 3Diii) of lipid droplets (LDs) in DIO+Fruc mice. These cardiomyocyte-accumulated LDs implied that fatty acid metabolism was dysregulated by excessive fructose intake.

**Figure 3.**
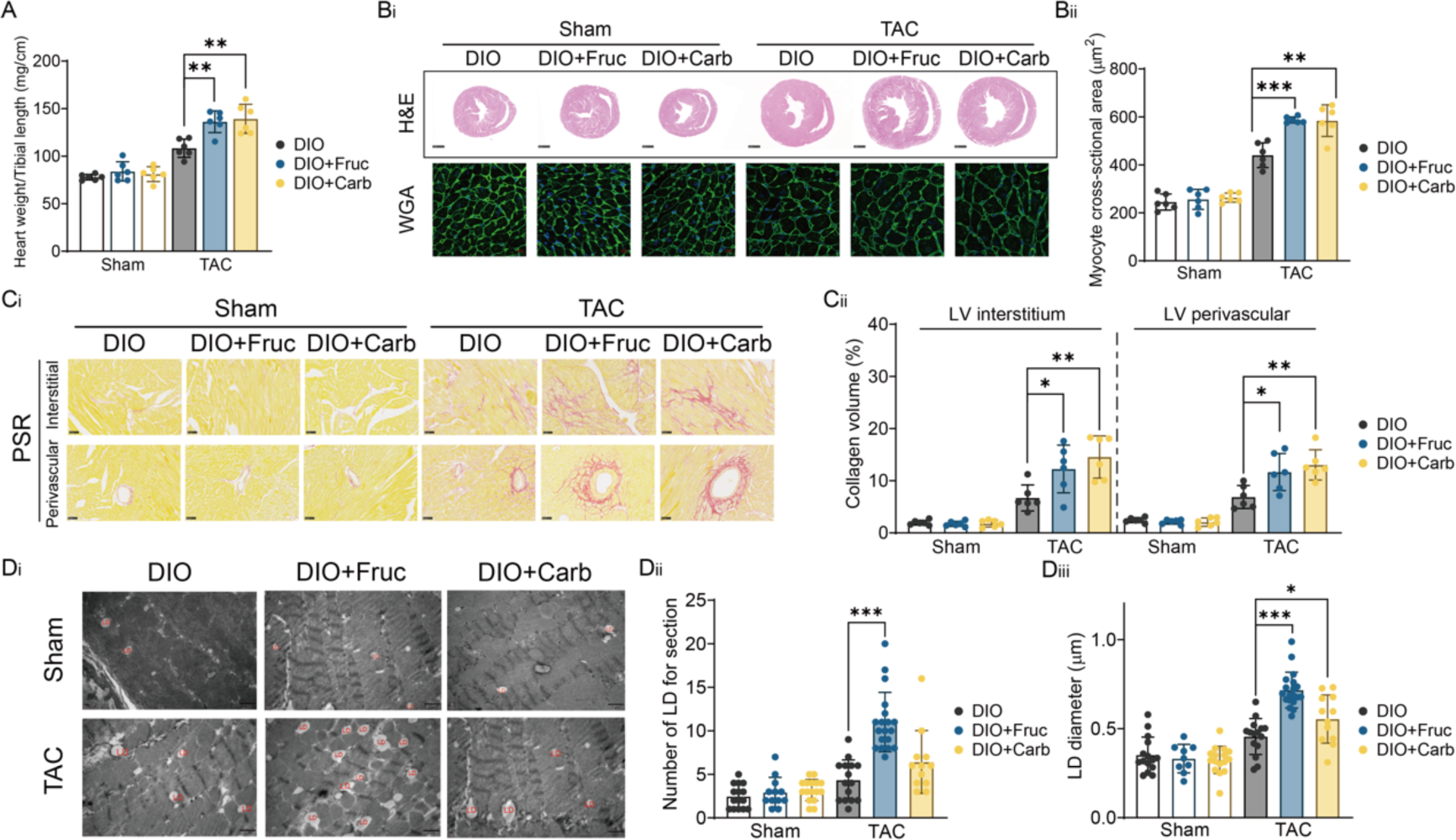
Excessive fructose exacerbated heart morphological lesions in DIO mice with transverse aortic constriction (TAC) challenge. (A) Heart weight to tibial length ratio was increased with TAC surgery, and DIO mice fed additional sugar (DIO+Carb) or drinking fructose (DIO+Fruc) were larger than regular DIO mice (\**p* < 0.05, \*\**p* < 0.01, n=6). (B) Both the cross-section of the heart (H&E staining) and cardiomyocytes (WGA staining) of DIO+Carb and DIO+Fruc mice hearts were larger than the regular DIO mouse. (C) PSR staining indicated that the collagen volume of DIO+Carb and DIO+Fruc mice was increased both interstitially and perivascularly following TAC challenging (*vs.* DIO, \**p* < 0.05, \*\**p* < 0.01, n=6). (D) Electron microscopy results indicated an accumulation number and enlarged volume of lipid droplets (LDs) in DIO+Fruc mouse (*vs.* DIO, \**p* < 0.05, \*\**p* < 0.01, \*\*\**p* < 0.001, n=6).

### Stable isotope labeling-based metabolic flux analysis indicates the suppression of heart fatty acid oxidation by excessive dietary fructose intake

While we identified LD accumulation in mouse heart tissue with excessive fructose administration, it was difficult to determine how and where these accumulated cardiac lipids originated: (1) *de novo* synthesized by the liver and delivered to the heart, or (2) synthesized in the heart, or (3) lipid surplus due to oxidation being suppressed. Thus, we designed an *in vivo* double stable isotopic labeling (DSIL) experiment to monitor the free fatty acid and fructose metabolism in mouse hearts simultaneously (Figure 4A). Oleic acid-d2 (OA-d2) and ^13^C6-fructose bolus were injected to trace fatty acid and fructose metabolic flux following three-week DIO or DIO+Fruc induction, and mouse tissues were harvested from 0 minutes to 30 minutes after the double stable isotopomers were administered. To comprehend fatty acid and fructose metabolic flux inside heart tissue, we evaluated isotopic labeling patterns of short-chain acyl-CoAs as these metabolites were strongly linked to fatty acid oxidation and fructose catabolism and would likely not export out of heart tissue. During beta-oxidation, the OA could constantly produce acetyl-CoAs alongside energy generation. Thus, the rate of acetyl-CoA, acetoacetyl-CoA, and 3-hydroxy-3-methylglutary-CoA (HMG-CoA) yield by OA-d2 was largely reduced (Figure 4Bi-Biv), suggesting a downregulation of fatty acid oxidation (FAO) in DIO+Fruc mouse hearts. In contrast, the higher labeling rate of glycerol-3-phosphate, lactate, and acetyl-CoA but lower labeling rate of pyruvate by ^13^C6-fructose suggested an elevation of fructolysis and fructose oxidation rate (Figure 4Ci-Cv) in excessive fructose-treated mouse hearts. Given that inhibition of either *de novo* synthesized fatty acids or lipids delivered from the liver to the heart could result in an *in situ* cardiac FAO downregulation, we monitored the isotopic labeled lipids and fructose in the liver and peripheral blood (PB) to understand fatty acid influx with additional fructose (Figure S5A). The assessment of both hepatic efflux and PB influx of ^13^C-fructose labeled total triacylglycerol (TG, Figure S5Bii-Biii) and diacylglycerol (DG, Figure S5Biv-Bv) were not altered under the administration of excessive fructose although the hepatic lipogenesis gene was slightly upregulated without any associated hepatic inflammation (Figure S5C-D). Similar to our metabolic flux analysis, the transcriptomic analysis of mouse heart tissues also indicated downregulation of cardiac fatty acid catabolic processes and FAO metabolism compared to the regular DIO mouse (Figure 4Di and Figure S6). Most FAO metabolism-associated genes, including *Cpt1a*, *Cpt1b*, *Fgf21*, and *Ppara*, were downregulated as expected (Figure 4Dii). Notably, a well-known carnitine palmitoyltransferase 1 (CPT1) inhibitor^32^, malonyl-CoA, was increased in DIO+Fruc mice (Figure 4Diii), which could directly suppress the CPT1-mediated FAO in the heart. These results indicated that excessive dietary fructose could directly suppress FAO in DIO mouse hearts other than fatty acid supply inhibition.

**Figure 4.**
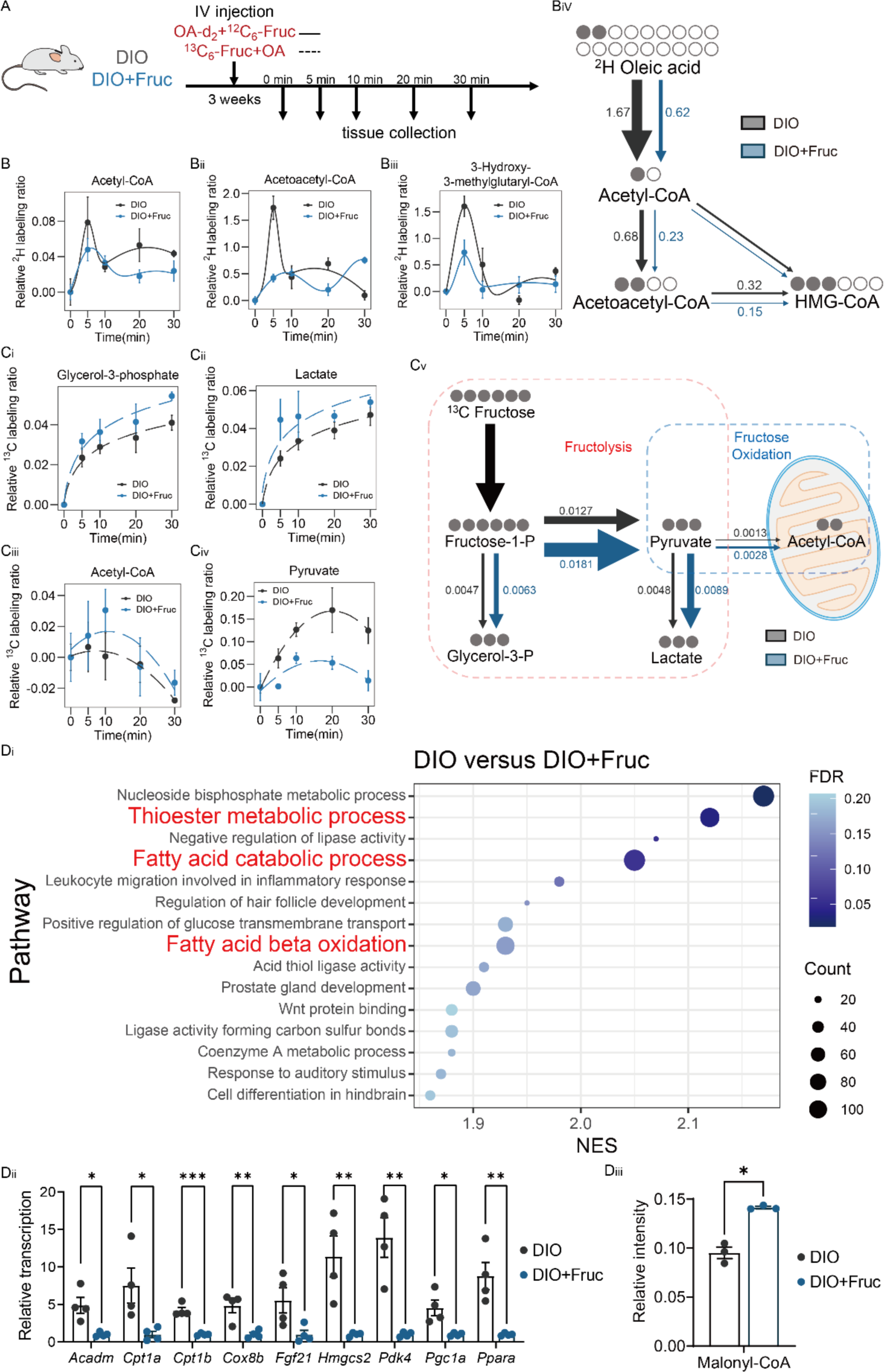
*In vivo* double stable isotopic labeling (DSIL) experiments suggested fatty acid oxidation suppression and fructolysis promotion in DIO+Fruc mice. (A) The workflow of DIO and DIO+Fruc mouse groups was isotopically labeled by oleic acid-d2 and ^13^C6-fructose. (B) The tracing of ^2^H labeling of (Bi) acetyl-CoA, (Bii) acetoacetyl-CoA, and (Biii) 3-hydroxy-3-methylglutaryl-CoA (HMG-CoA) flux decrease indicated (Biv) oleic acid oxidation was suppressed in DIO+Fruc mouse (n=3 in each group and time point). (C) The tracing of ^13^C labeling of (Ci) glycerol-3-phosphate, (Cii) lactate, and (Ciii) acetyl-CoA flux increase and (Civ) pyruvate flux decrease indicated (Cv) fructolysis was promoted in DIO+Fruc mouse (n=3 in each group and time point). (Di) RNA-sequencing and (Dii) real-time PCR results indicated downregulation of FAO association gene expression in DIO+Fruc mice (*vs.* DIO, \**p* < 0.05, \*\**p* < 0.01, \*\*\**p* < 0.001, n=4), and endogenous CPT1B inhibitor (Diii) malonyl-CoA level was increased in DIO+Fruc mice (*vs.* DIO, \**p* < 0.05, n=3).

### Excessive fructose intake suppresses fatty acid oxidation via the AMPK-ACC axis

As CPT1 activity was mostly suppressed by acetyl coenzyme A carboxylase 1/2 (ACC1/2)-induced malonyl-CoA^27^, it is possible that excessive fructose could regulate fatty acid metabolism via ACC activity alteration^28,29^. We investigated the phosphoproteome and found phosphorylated ACC2 levels were downregulated in DIO+Fruc mouse heart tissues (Figure 5Ai-Aii). Interestingly, some other AMPK-associated downstream substrates, including 3-hydroxy-3-methylglutaryl-CoA reductase (HMGCR), were also highly phosphorylated^28^ (Figure 5Ai and S7A-B), while an endogenous AMPK activator, 5-aminoimidazole-4-carboxamide ribonucleotide (AICAR), was elevated (Figure 5B). These findings suggested that excessive fructose might also limit cardiac FAO metabolism through AMPK-ACC axis mediation. As AMPK and phosphorylated AMPK were altered rapidly *ex vivo*, we harvested fresh heart tissues of DIO and DIO+Fruc mice either fed *ad libitum* or starved. As anticipated, the p-AMPK/AMPK ratio was significantly reduced (Figure 5 Ci-Cii) in excessive fructose-fed *ad libitum* mouse hearts, which was accompanied by decreased phosphorylated ACC (Figure 5Ci and Ciii-Cv). Additionally, the levels of CPT1-α and CPT1-β were also downregulated in extra fructose-fed *ad libitum* mouse hearts and continued to decrease following a 6-hour starvation (Figure 5Di-Diii). As the *Acc2* gene could also be regulated by another fructose-regulated transcriptional factor, *Chrebp*^30,31^, we evaluated the downstream gene targets of *Chrebp* but did not identify substantial changes (Figure S7C). This indicated that the expression of ACC2 was not associated with *Chrebp*. These findings suggested that excessive fructose in the heart competed for fatty acids for oxidation metabolism and inhibited FAO through AMPK-ACC axis mediation.

**Figure 5.**
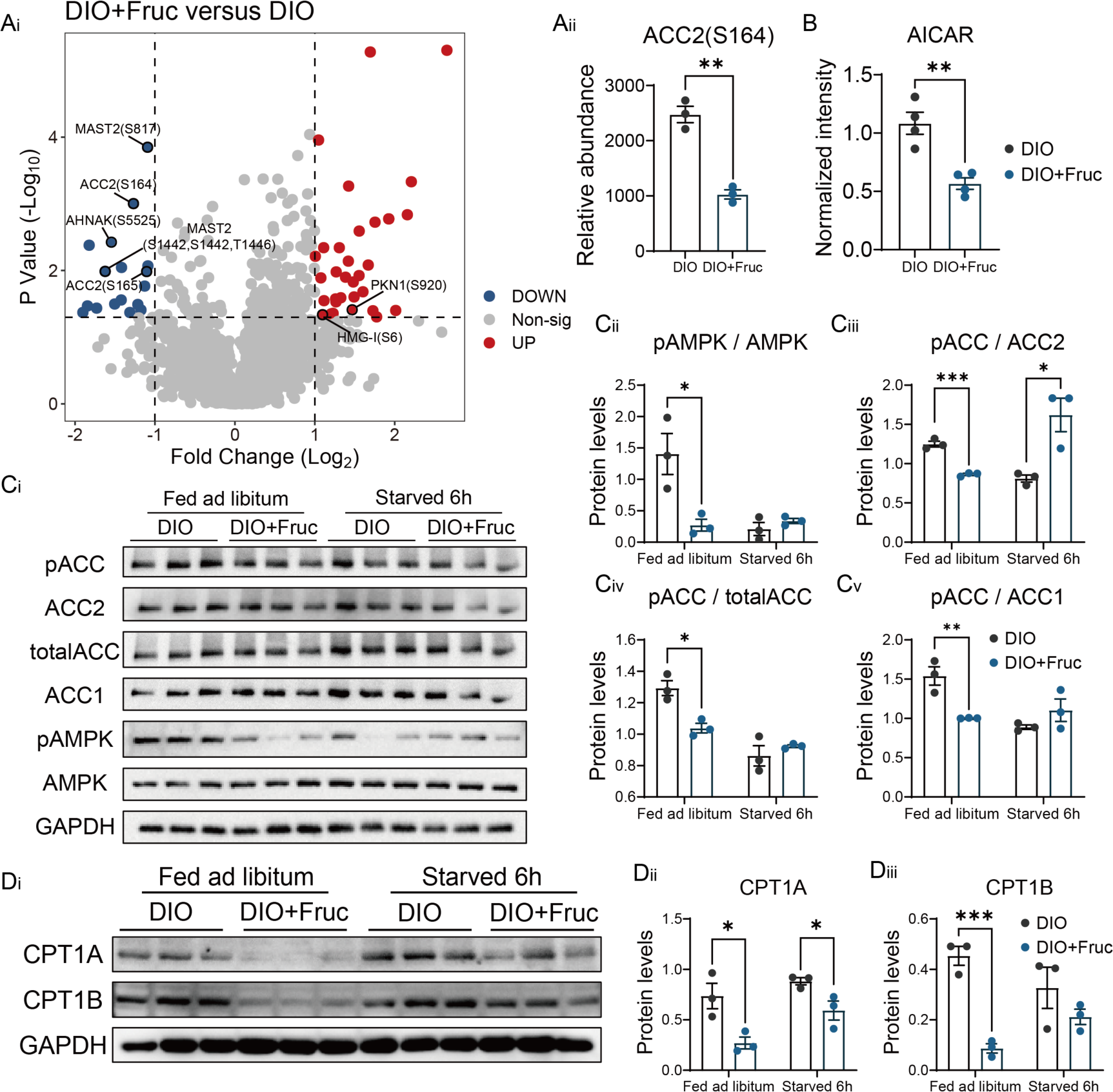
Excessive fructose could suppress ACC and AMPK phosphorylation levels. (A) The phosphoproteome results indicated downregulation of ACC2 and other AMPK-associated substrate phosphorylation levels in DIO+Fruc mouse heart (*vs.* DIO, \**p* < 0.05, \*\**p* < 0.01, n=3); (B) The endogenous AMPK activator AICAR was upregulated in DIO+Fruc mouse hearts (*vs.* DIO, \**p* < 0.05, \*\**p* < 0.01, n=3). (C) The phosphorylation of AMPK and ACC were downregulated when fed *ad libitum*, but unaltered following a 6-hour fast in the fresh DIO+Fruc mouse heart tissue (*vs.* DIO, \**p* < 0.05, n=3); while (D) CPT1-α and CPT1-β levels were downregulated fed ad libitum and after 6-hour fasting in fresh DIO+Fruc mouse heart tissue (*vs.* DIO, \**p* < 0.05, \*\**p* < 0.01, n=3).

### Activating AMPK or deactivating ACC attenuates the excessive fructose intake-induced heart function impairment in DIO mice

To further identify whether the fructose-induced cardiac FAO suppression could be mediated through the AMPK-ACC axis, we then treated excessive fructose-fed, and TAC-challenged DIO mice with either AMPK activator (AICAR) or ACC inhibitor (CP-640186) and assessed their heart function alterations (Figure 6A). Upon TAC challenge, the echocardiographic results indicated that DIO+Fruc mice had impaired diastolic (E/E’) and systolic function (LVEF, FS) as anticipated. However, such cardiac impairments were significantly recovered after AICAR or CP-640186 treatments (Figure 6Bi-Biii). Notably, even in the sham group, mice treated with ACC inhibitor had improved diastolic and systolic function (Figure 6Bi-Biii). Similarly, the histological results of mice hearts indicated enlargement of both total heart tissue (Figure 6Ci, H&E staining) and cross-sectional cardiomyocyte area (Figure 6Ci-Cii, WGA staining). This could be observed in DIO+Fruc mice, but activation of AMPK or inhibition of ACC could limit such enlarged myocardial hypertrophy (Figure 6Ci-Cii). PSR staining of heart tissue demonstrated that the enlarged interstitial and perivascular fibrosis in DIO+Fruc mice upon TAC challenge was improved with AICAR or CP-640186 treatment (Figure 6Di-Dii). Additionally, AICAR or CP-640186 treatment can slightly downregulate the thickened LVPW;d in DIO+Fruc mouse, but not significantly (Figure S8A-B). These findings indicated that the activation of AMPK or deactivation of ACC could attenuate the cardiac impairment produced by excessive fructose treatment in DIO mice.

**Figure 6.**
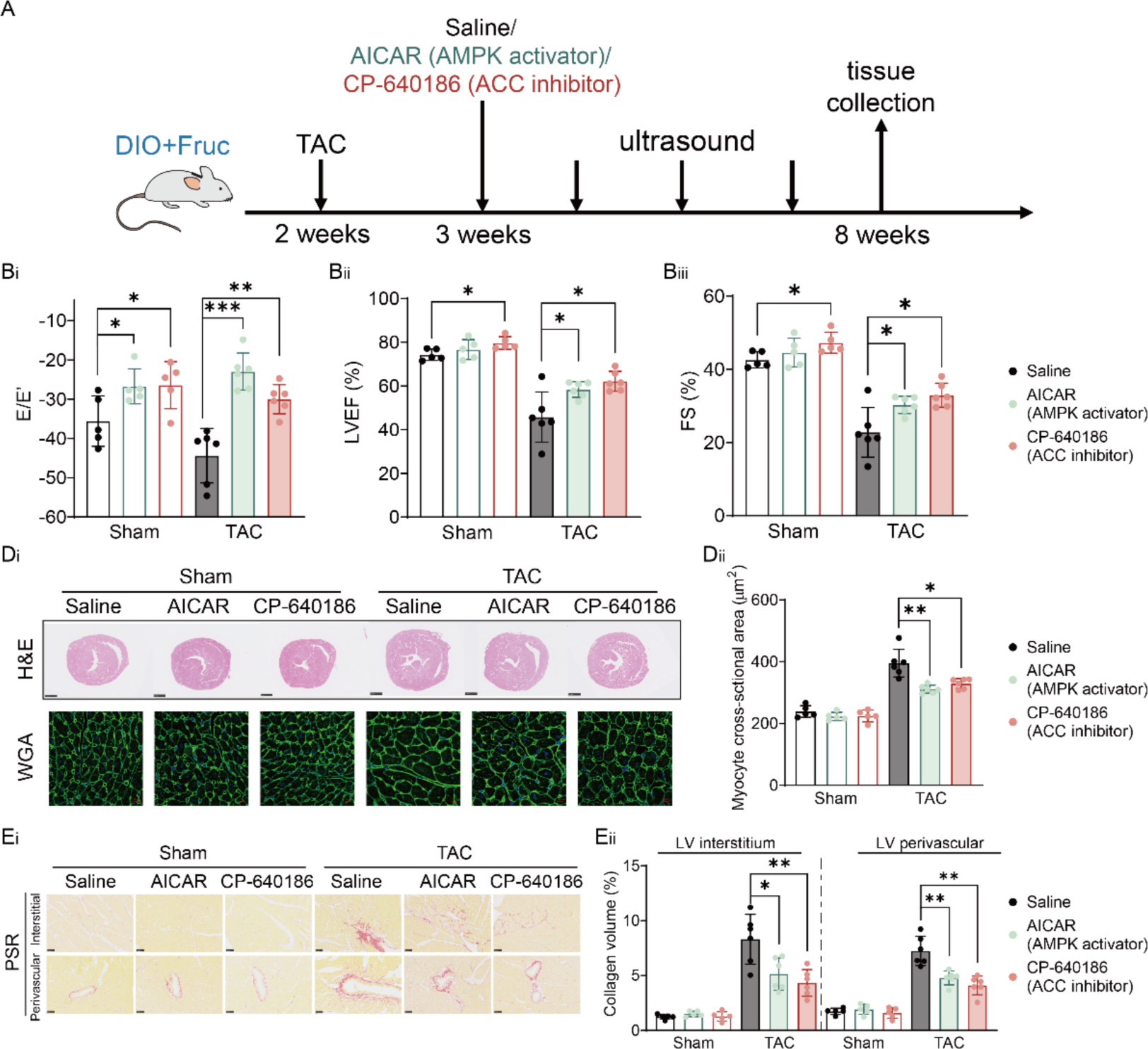
Chemical activation of AMPK or deactivation of ACC attenuates cardiac impairment induced by excessive fructose treatment in DIO mice. (A) Workflow of DIO+Fruc mouse drug treatments. The declined (Bi) Left ventricular ejection fractions (LVEF), (Bii) fractional shortening (FS), and (Biii) E/E’ following TAC surgery in DIO+Fruc mice were improved with AICAR or CP-640186 treatment, respectively. (vs. Saline, \**p* < 0.05, \*\**p* < 0.01, \*\*\**p* < 0.001, n=6). Both the (Ci) cross-section of the heart (H&E staining) and (Ci-Cii) cardiomyocytes (WGA staining) enlargements of DIO+Fruc mice hearts were shrunk upon AICAR or CP-640186 treatment, respectively. (vs. Saline, \**p* < 0.05, \*\**p* < 0.01, n=6). (D) The PSR staining indicated that elevated collagen volume of DIO+Fruc mice was decreased with AICAR or CP-640186 treatment both interstitially and perivascularly upon TAC challenge, respectively. (*vs.* Saline, \**p*<0.05, \*\**p*<0.01, n=6)

### High blood fructose levels are associated with impaired heart function in aortic stenosis patients

As excessive fructose intake could induce heart dysfunctions in mice, it was necessary to examine if excessive fructose levels were also associated with clinical heart dysfunction. We obtained 27 blood samples of aortic stenosis patients (Table S1) and measured the plasma fructose level. As anticipated, the fructose level instead of glucose or sucrose (Figure S9A-D) was significantly correlated to ejection fraction (EF) decline (Figure 7A) and fractional shortening (Figure 7B). This suggested that the plasma level of fructose was highly correlated to clinical heart dysfunction.

**Figure 7.**
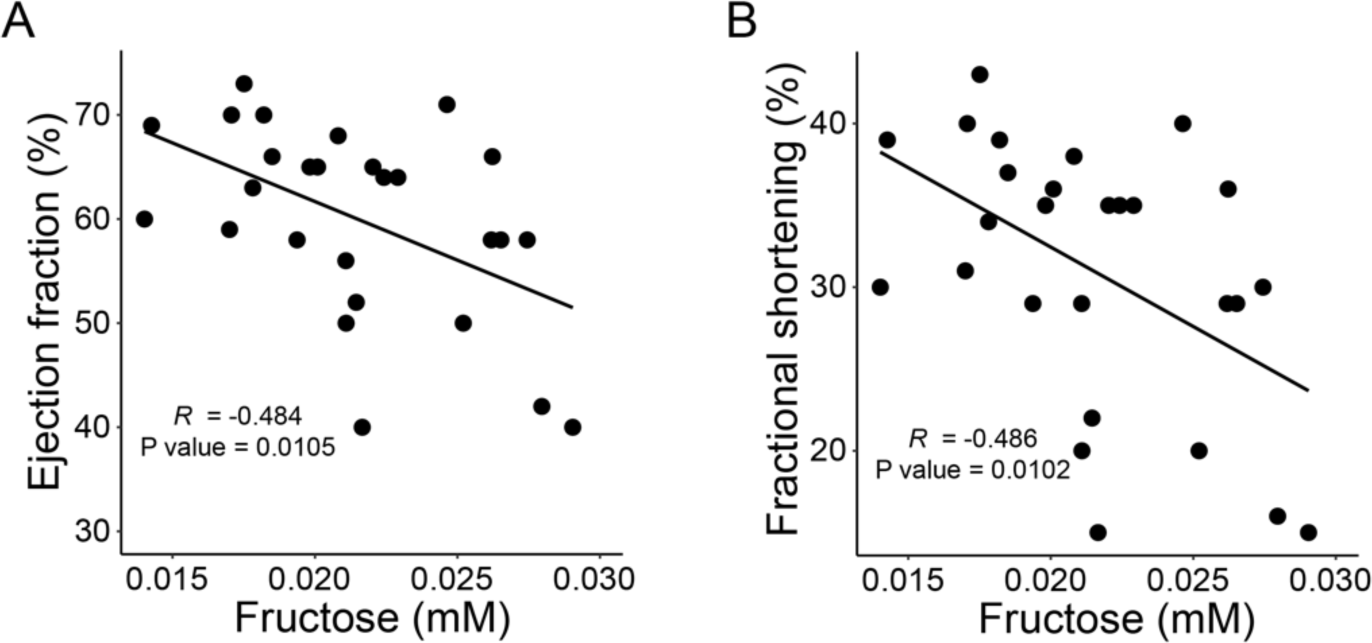
High fructose levels were associated with heart dysfunction in aortic stenosis patients. (A) the ejection fraction and (B) fractional shortening were reduced in aortic stenosis patients with a higher concentration of plasma fructose (n=27).

## Discussion

In this study, we identified that the intake of excessive fructose could alter cardiac lipid metabolism and inhibit FAO in a DIO mouse model, leading to systolic dysfunction. A recent meta-analysis also indicated that drinking high-sugar beverages could correlate to a high risk of cardiovascular diseases, type 2 diabetes, as well as liver cancers^32–35^. It pointed out that the abuse of added sugar worldwide should be carefully considered. Except for artificial sweeteners, high-fructose corn syrup (HFCS) is one of the most popular added sugars in modern industrialized foods and beverages^36^. Although fructose and glucose are very similar isomers, the uptake and metabolism rate of fructose is much higher than that of glucose due to the absence of phosphofructokinase (PFK) inhibition in fructolysis^36^. Therefore, the rapid absorption and metabolic rate of dietary HFCS or fructose might be the primary cause of such heart metabolic dysregulation. Our *in vivo* double stable isotopic tracing experiment clearly demonstrated that the FAO metabolic flux was inhibited by rapid competition for oxygen from fructose catabolism. Additionally, excessive fructose could also occupy the fructolysis pathway to generate fewer ATPs than the equivalent moles of fatty acid combustion. Mirtschink *et al.* found that enforced fructolysis could lead to mouse heart disease as well^37^. We believed that energy supplementary deficiency involved with excessive fructose intake could be the primary cause of such heart failure.

Maintaining metabolic balance in the myocardium is a primary therapeutic approach for heart failure, and fatty acids operate as the predominant substrates for ATP production (> 70%) in the heart^10^. The prohibition of FAO metabolism would cause heart dysfunction or further adverse outcomes. Shao *et al.* found that increasing 40% FAO by deletion of ACC2 could abolish cardiomyopathy, which could rescue mitochondrial function and restore cardiac function in prolonged high-fat diet-fed mice^38^. Similarly, our findings indicated the chemical inhibition of ACC could also attenuate heart failure and myocardial hypertrophy. These outcomes implied that the promotion of FAO for energy production could benefit heart function.

However, whether to select inhibition or stimulation of FAO as therapy for patients with heart failure is still debatable^39^. Perhexiline and etomoxir were mainly targeted on the inhibition of FAO by suppressing CPT1 activity, and trimetazidine was aimed at long-chain 3-ketoacyl-CoA thiolase (KAT) inhibition^39^. Moreover, Xiang *et al.* observed that the inhibition of fatty acid oxidation via inactivating *Cpt1b* could produce heart regeneration in adult mice^9^. Such controversial observations were most likely due to cardiac energy generation and oxygen consumption balance at the stage of heart failing. Given that the oxidation of fatty acid processes could generate abundant ATP but consume a correspondingly high amount of oxygen^40^, the use of fatty acids as an essential energy source was quite unique for the adult mammalian heart. A healthy heart can readily obtain sufficient oxygens for FAO energy generation, but a failing heart might lack the ability to oxidize all the fatty acids and result in lipid accretion. Thus, the inhibition of FAO could limit such oxygen consumption stress and promote other anaerobic catabolic mechanisms such as fructolysis and glycolysis. However, the promotion of FAO (by deactivation of ACC) could provide more ATPs and eliminate the LD accumulation at a relatively earlier time point.

As an important energy-sensing molecule, AMPK exerts an important role in this excessive fructose-mediated FAO regulation. Our findings implied that fructose could also be sensed by AMPK, and supplementation of exogenous AICAR to activate AMPK could attenuate the heart failure phenotype induced by excessive fructose treatment. Some animal studies have indicated that exogenous treatment of AICAR has beneficial effects on failing hearts^41,42^. Chen-Song *et al.* demonstrated that AMPK could sense glucose via aldolase A^29,43^, and we suspected that AMPK could also sense the concentration of fructose via aldolase through aldolase B (ALDOB). The upregulation of *Aldob* gene expression was observed in excessive fructose-treated mouse heart tissues, as expected (Figure S10). More sophisticated biochemical demonstrations have been planned in our future experiments.

A limitation of the study is the lack of access to the clinical measurement of fructose intake. The correlation of fructose level and patient heart function was based on the measurement of plasma fructose level. Moreover, a clinically stable isotopic tracing experiment would be ideal to validate the results of excessive fructose-caused heart metabolism alterations.

## Conclusions

In summary, our findings indicate that excessive dietary fructose could induce heart failure by suppressing cardiac FAO metabolism in DIO mice. Our findings suggest that limiting dietary fructose intake, especially HFCS-sweetened soft beverages, could prevent such outcomes and medically promote heart FAO metabolism, which might restore impaired heart function therapeutically.

## Acknowledgments

We thank the Single Cell Quantitative Metabolomics and Lipidomics Core Facility of IMIB at Fudan University for generating metabolomic and lipidomic data.

## References

1. Popkin, B. M., Adair, L. S. & Ng, S. W. Global nutrition transition and the pandemic of obesity in developing countries. Nutr. Rev. 70, 3–21 (2012).

2. Castro, J. P., El-Atat, F. A., McFarlane, S. I., Aneja, A. & Sowers, J. R. Cardiometabolic syndrome: pathophysiology and treatment. Curr. Hypertens. Rep. 5, 393–401 (2003).

3. Alberti, K. G. M. M. et al. Harmonizing the metabolic syndrome: a joint interim statement of the International Diabetes Federation Task Force on Epidemiology and Prevention; National Heart, Lung, and Blood Institute; American Heart Association; World Heart Federation; International. Circulation 120, 1640–1645 (2009).

4. Bray, G. A., Nielsen, S. J. & Popkin, B. M. Consumption of high-fructose corn syrup in beverages may play a role in the epidemic of obesity. Am. J. Clin. Nutr. 79, 537–543 (2004).

5. Malik, V. S., Popkin, B. M., Bray, G. A., Després, J.-P. & Hu, F. B. Sugar-sweetened beverages, obesity, type 2 diabetes mellitus, and cardiovascular disease risk. Circulation 121, 1356–1364 (2010).

6. Yang, Q. et al. Added sugar intake and cardiovascular diseases mortality among US adults. JAMA Intern. Med. 174, 516–524 (2014).

7. de Koning, L. et al. Sweetened beverage consumption, incident coronary heart disease, and biomarkers of risk in men. Circulation 125, 1735–41, S1 (2012).

8. Horton, J. L., et al. The failing heart utilizes 3-hydroxybutyrate as a metabolic stress defense. JCI insight 4, (2019).

9. Li, X. et al. Inhibition of fatty acid oxidation enables heart regeneration in adult mice. Nature 622, 619–626 (2023).

10. Lopaschuk, G. D., Karwi, Q. G., Tian, R., Wende, A. R. & Abel, E. D. Cardiac Energy Metabolism in Heart Failure. Circ. Res. 128, 1487–1513 (2021).

11. Taegtmeyer, H. & Overturf, M. L. Effects of moderate hypertension on cardiac function and metabolism in the rabbit. *Hypertens. (Dallas*, Tex. 1979) 11, 416–426 (1988).

12. Sack, M. N. et al. Fatty acid oxidation enzyme gene expression is downregulated in the failing heart. Circulation 94, 2837–2842 (1996).

13. Stanley, W. C., Recchia, F. A. & Lopaschuk, G. D. Myocardial substrate metabolism in the normal and failing heart. Physiol. Rev. 85, 1093–1129 (2005).

14. Dávila-Román, V. G. et al. Altered myocardial fatty acid and glucose metabolism in idiopathic dilated cardiomyopathy. J. Am. Coll. Cardiol. 40, 271–277 (2002).

15. Neglia, D. et al. Impaired myocardial metabolic reserve and substrate selection flexibility during stress in patients with idiopathic dilated cardiomyopathy. Am. J. Physiol. Heart Circ. Physiol. 293, H3270–8 (2007).

16. Ahluwalia, N., Dwyer, J., Terry, A., Moshfegh, A. & Johnson, C. Update on NHANES Dietary Data: Focus on Collection, Release, Analytical Considerations, and Uses to Inform Public Policy. Adv. Nutr. 7, 121–134 (2016).

17. Rhodes, D. G., Morton, S., Myrowitz, R. & Moshfegh, A. J. Food and Nutrient Database for Dietary Studies 2019–2020: An application database for national dietary surveillance. J. Food Compos. Anal. 123, 105547 (2023).

18. Shin, S.-Y. et al. An atlas of genetic influences on human blood metabolites. Nat. Genet. 46, 543–550 (2014).

19. Long, T. et al. Whole-genome sequencing identifies common-to-rare variants associated with human blood metabolites. Nat. Genet. 49, 568–578 (2017).

20. van der Harst, P. & Verweij, N. Identification of 64 Novel Genetic Loci Provides an Expanded View on the Genetic Architecture of Coronary Artery Disease. Circ. Res. 122, 433–443 (2018).

21. Wojcik, G. L. et al. Genetic analyses of diverse populations improves discovery for complex traits. Nature 570, 514–518 (2019).

22. Shah, S. et al. Genome-wide association and Mendelian randomisation analysis provide insights into the pathogenesis of heart failure. Nat. Commun. 11, 163 (2020).

23. Elsworth, B. et al. The MRC IEU OpenGWAS data infrastructure. bioRxiv 2020.08.10.244293 (2020). doi:10.1101/2020.08.10.244293

24. Hemani, G. et al. The MR-Base platform supports systematic causal inference across the human phenome. Elife 7, (2018).

25. Zhang, X. et al. Metabolic disorder in the progression of heart failure. Sci. China. Life Sci. 62, 1153–1167 (2019).

26. Heinonen, I. et al. The effects of equal caloric high fat and western diet on metabolic syndrome, oxidative stress and vascular endothelial function in mice. Acta Physiol. (Oxf). 211, 515–527 (2014).

27. Foster, D. W. Malonyl-CoA: the regulator of fatty acid synthesis and oxidation. J. Clin. Invest. 122, 1958–1959 (2012).

28. Chen, Z. et al. Global phosphoproteomic analysis reveals ARMC10 as an AMPK substrate that regulates mitochondrial dynamics. Nat. Commun. 10, 104 (2019).

29. Zhang, C.-S. et al. Fructose-1,6-bisphosphate and aldolase mediate glucose sensing by AMPK. Nature 548, 112–116 (2017).

30. Shi, J.-H. et al. Fructose overconsumption impairs hepatic manganese homeostasis and ammonia disposal. Nat. Commun. 14, 7934 (2023).

31. Shi, J.-H. et al. Liver ChREBP Protects Against Fructose-Induced Glycogenic Hepatotoxicity by Regulating L-Type Pyruvate Kinase. Diabetes 69, 591–602 (2020).

32. Witkowski, M. et al. The artificial sweetener erythritol and cardiovascular event risk. Nat. Med. 29, 710–718 (2023).

33. Debras, C. et al. Artificial Sweeteners and Risk of Type 2 Diabetes in the Prospective NutriNet-Santé Cohort. Diabetes Care (2023). doi:10.2337/dc23-0206

34. Debras, C. et al. Artificial sweeteners and risk of cardiovascular diseases: results from the prospective NutriNet-Santé cohort. BMJ 378, e071204 (2022).

35. Zhao, L. et al. Sugar-Sweetened and Artificially Sweetened Beverages and Risk of Liver Cancer and Chronic Liver Disease Mortality. JAMA 330, 537–546 (2023).

36. Mirtschink, P., Jang, C., Arany, Z. & Krek, W. Fructose metabolism, cardiometabolic risk, and the epidemic of coronary artery disease. Eur. Heart J. 39, 2497–2505 (2018).

37. Mirtschink, P. et al. HIF-driven SF3B1 induces KHK-C to enforce fructolysis and heart disease. Nature 522, 444–449 (2015).

38. Shao, D. et al. Increasing Fatty Acid Oxidation Prevents High-Fat Diet–Induced Cardiomyopathy Through Regulating Parkin-Mediated Mitophagy. Circulation 142, 983–997 (2020).

39. Ritterhoff, J. & Tian, R. Metabolic mechanisms in physiological and pathological cardiac hypertrophy: new paradigms and challenges. Nat. Rev. Cardiol. 20, 812–829 (2023).

40. Schönfeld, P. & Reiser, G. Why does brain metabolism not favor burning of fatty acids to provide energy? Reflections on disadvantages of the use of free fatty acids as fuel for brain. J. Cereb. blood flow Metab. Off. J. Int. Soc. Cereb. Blood Flow Metab. 33, 1493–1499 (2013).

41. Cieslik, K. A. et al. AICAR-dependent AMPK activation improves scar formation in the aged heart in a murine model of reperfused myocardial infarction. J. Mol. Cell. Cardiol. 63, 26–36 (2013).

42. Li, Y. et al. AMPK blunts chronic heart failure by inhibiting autophagy. Biosci. Rep. 38, (2018).

43. Li, M. et al. Aldolase is a sensor for both low and high glucose, linking to AMPK and mTORC1. Cell research 31, 478–481 (2021).

