## Supplemental Figures for "Excessive Dietary Fructose Aggravates Heart Failure via Impairing Myocardial Fatty Acid Oxidation Metabolism in Diet Induced Obese Mouse"

1. Shanghai Key Laboratory of Metabolic Remodeling and Health, Institute of Metabolism and Integrative Biology and School of Life Sciences, Fudan University, Shanghai, 200438, China;
2. Department of Cardiology, Institute of Vascular Medicine, Peking University Third Hospital, NHC Key Laboratory of Cardiovascular Molecular Biology and Regulatory Peptides, State Key Laboratory of Vascular Homeostasis and Remodeling Peking University, Beijing, 100191, China;
3. School of Life Sciences, Inner Mongolia University, Hohhot Inner Mongolia, 010021, China;
4. Beijing Institute of Heart, Lung and Blood Vessel Diseases, Beijing Anzhen Hospital, Capital Medical University, Beijing 100029, China;
5. Department of Cardiology, Peking University First Hospital, Beijing, 100034, China;
6. Shanghai Institute of Nutrition and Health, Chinese Academy of Sciences, Shanghai, 200031, China;
7. Lead contact;

### These authors contribute equally

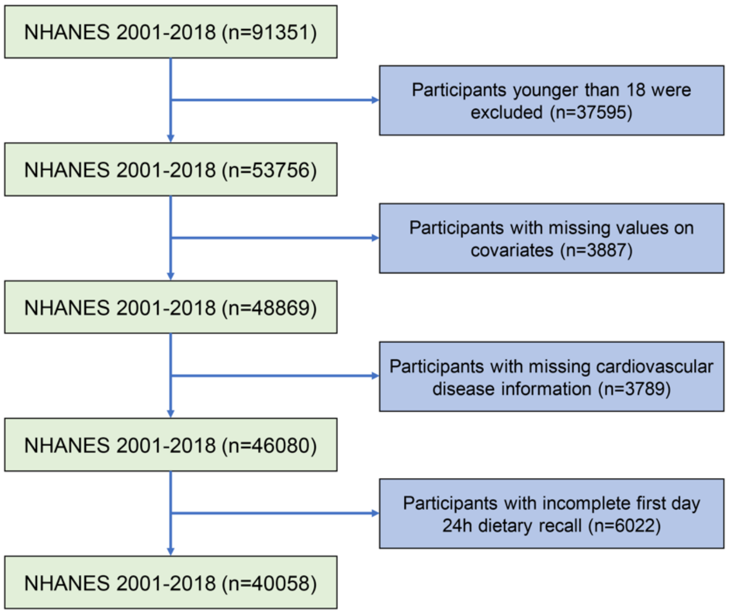


Figure S1. Workflow of meta-analysis for nutrient intake and association with cardiovascular diseases using National Health and Nutrition Examination Survey (NHANES).

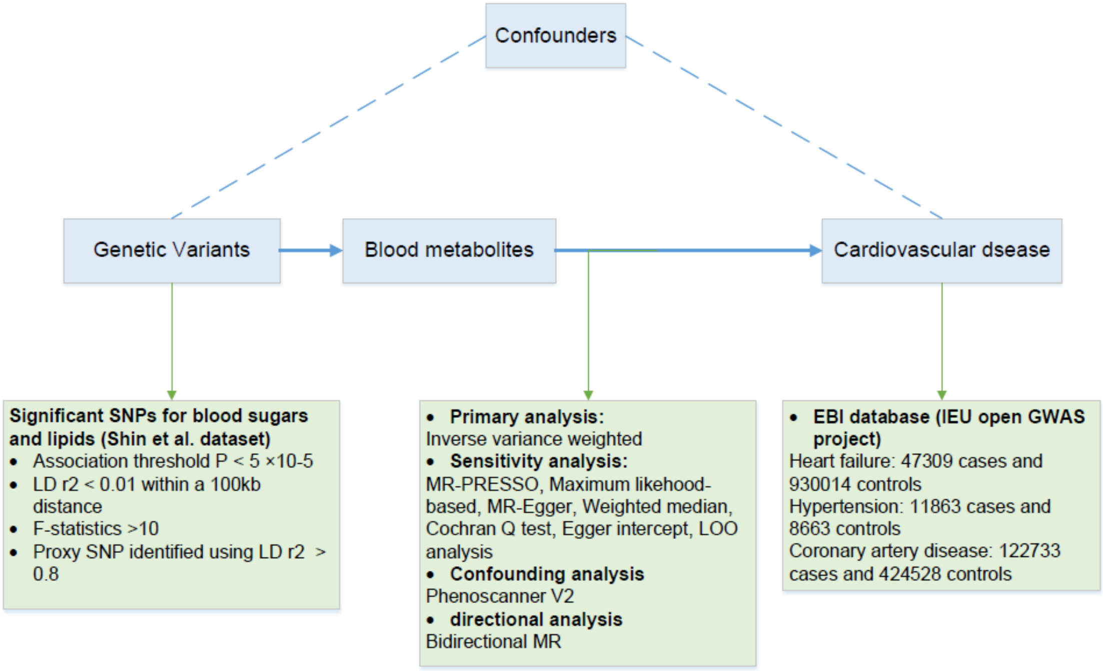


Figure S2. Workflow of two-sample Mendelian randomization analysis for three added sugars and causal association with cardiovascular diseases using published databases.

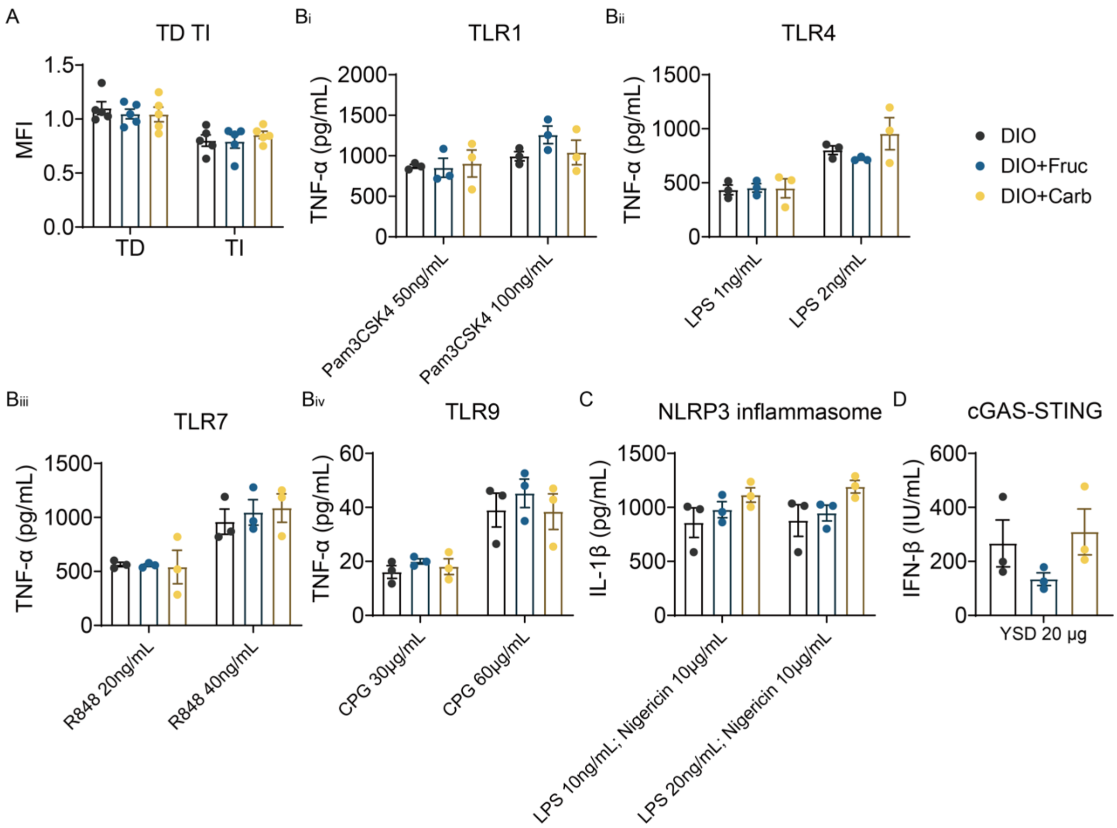


Figure S3. The immune responses were not changed after three-week fructose added DIO dietary treatments. (A) Thymus-dependent antigen (TD) and thymus-independent antigen (TI); (B) Toll-like receptors such as TLR1, TLR4, TLR7 and TLR9; and (C) interleukin-1β (IL-1β) and (D) interferon (IFN-β) responses were comparable to DIO treatment (*vs*. DIO, n=3).

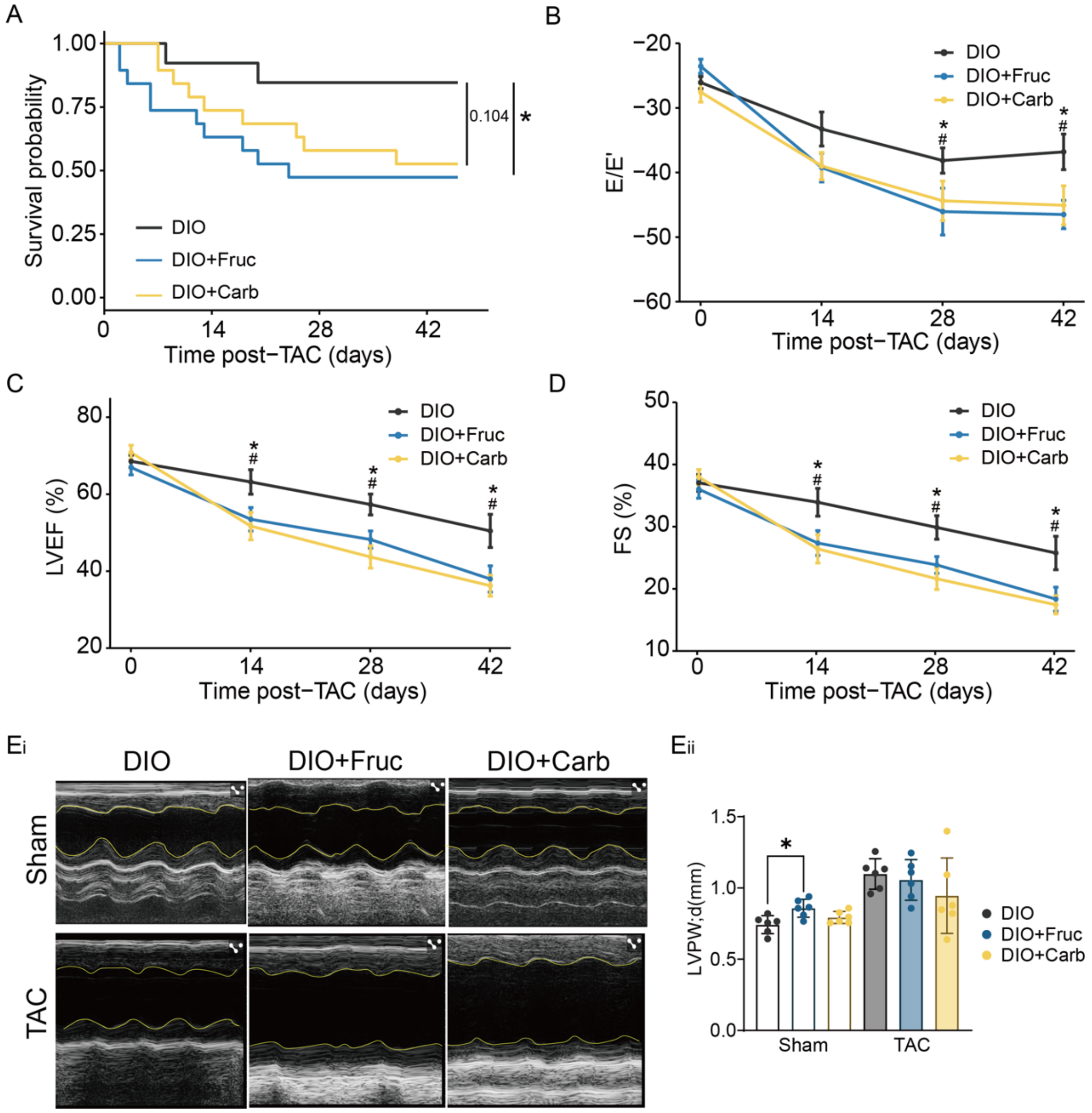


Figure S4. Excessive fructose impairs heart function in DIO mouse with transverse aortic constriction (TAC) challenging. (A) Survival probabilities were decreased in extra fructose fed mice (DIO+Fruc and DIO+Carb, *vs*. DIO, *p<0.05, **p<0.01, n=6). (B) E/E’, (C) Left ventricular ejection fractions (LVEF), and (D) fractional shortening (FS) were declined after TAC surgery in DIO mice fed with extra added sugar (DIO+Carb) or drinking fructose (DIO+Fruc); but (E) LVPW;d was not changed (*vs*. DIO, *p<0.05, **p<0.01, n=6).

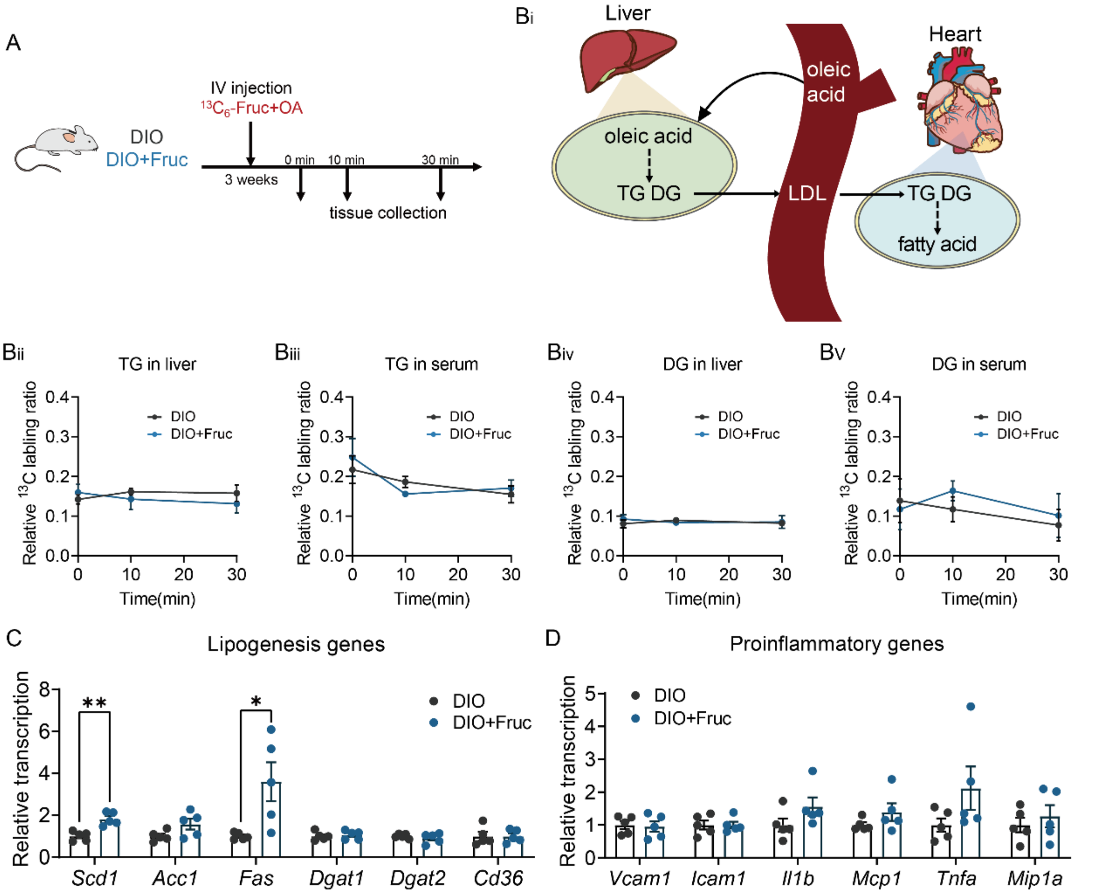


Figure S5. The *in vivo* stable isotopic labeling experiments indicated fatty acid delivery flux from liver did not change in DIO+Fruc mouse. (A) Workflow of DIO and DIO+Fruc mouse groups were isotopic labeled by oleic acid and $^{13}C_6$-fructose. ($B_i$) The pathway of oleic acid synthesized in liver and delivered to circulation system. The flux of triglycerides between ($B_{ii}$) liver and ($B_{iii}$) serum, and diglycerides between ($B_{iv}$) liver and ($B_v$) serum did not change in DIO+Fruc mouse (*vs.* DIO, n=3); The (C) hepatic lipogenesis genes were upregulated but not cause any (D) hepatic inflammations in DIO+Fruc mouse (*vs.* DIO, n=3).

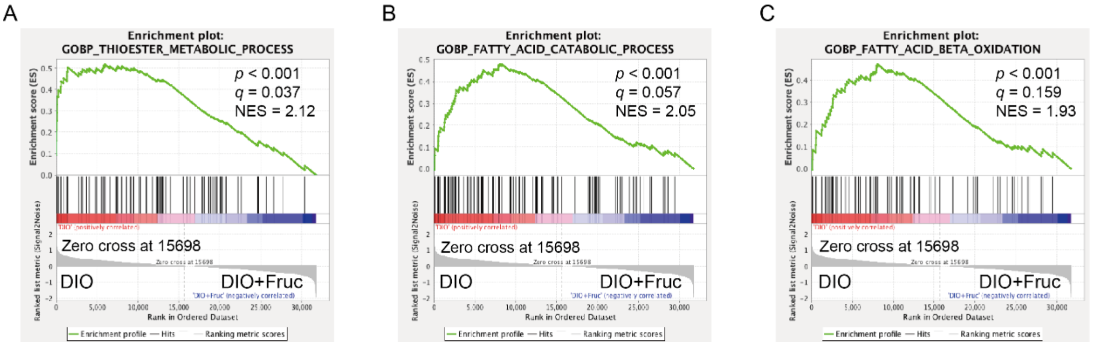


Figure S6. Gene set enrichment analysis (GSEA) results of DIO and DIO+Fruc mouse heart tissue transcriptome. The (A) thioester metabolic process, (B) fatty acid catabolic process, and (C) fatty acid beta oxidation pathways were downregulated in DIO+Fruc mouse heart.

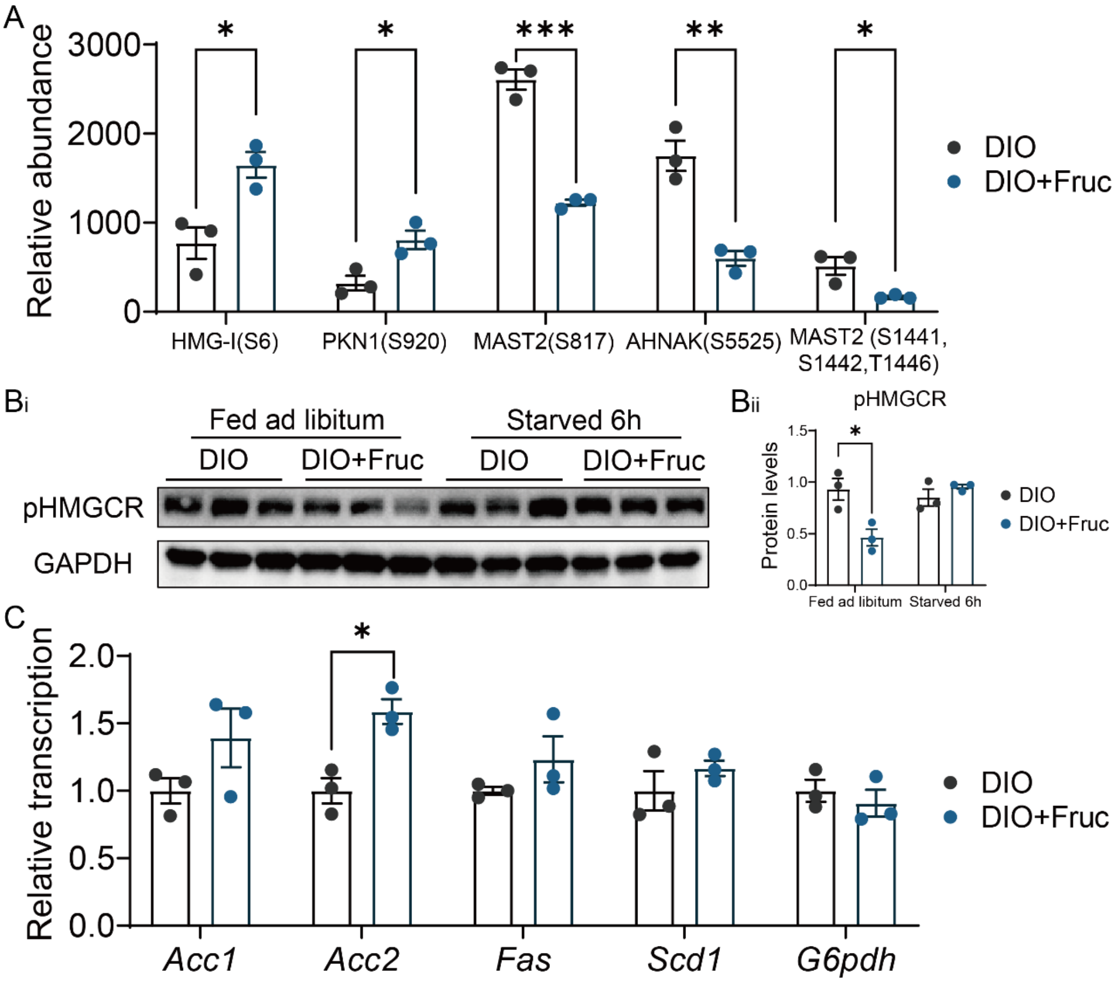


Figure S7. Excessive fructose could affect AMPK associated protein phosphorylation levels but not *Chrebp* downstream genes. (A) The phosphoproteome result indicated AMPK associated substrate phosphorylation levels were affected in DIO+Fruc mouse heart (vs. DIO, *p<0.05, **p<0.01, n=3); (B) The level of HMGCR was downregulated in DIO+Fruc mouse heart (vs. DIO, *p<0.05, **p<0.01, n=3); (C) The *Chrebp* downstream genes were not thoroughly upregulated except the *Acc2* in DIO+Fruc mouse heart (vs. DIO, *p<0.05, n=3).

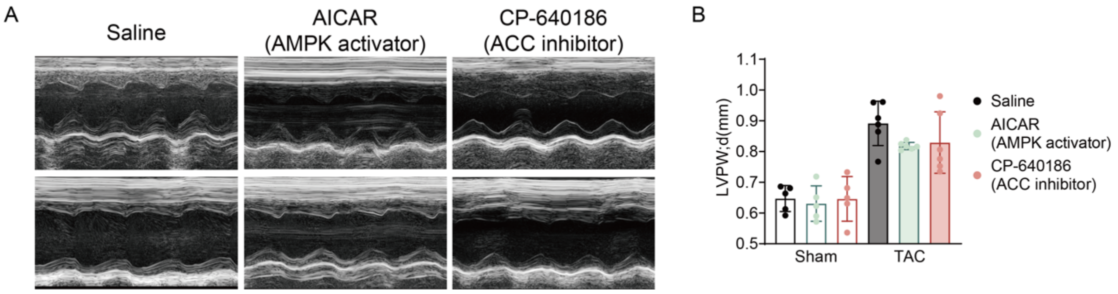


Figure S8. Chemical activation of AMPK or deactivation of ACC did not affect LVPW;d by excessive fructose treatment in DIO mouse. LVPW;d were comparable after TAC surgery in DIO+Fruc mice with AICAR or CP-640186 treatment, respectively. (*vs.* Saline, n=6).

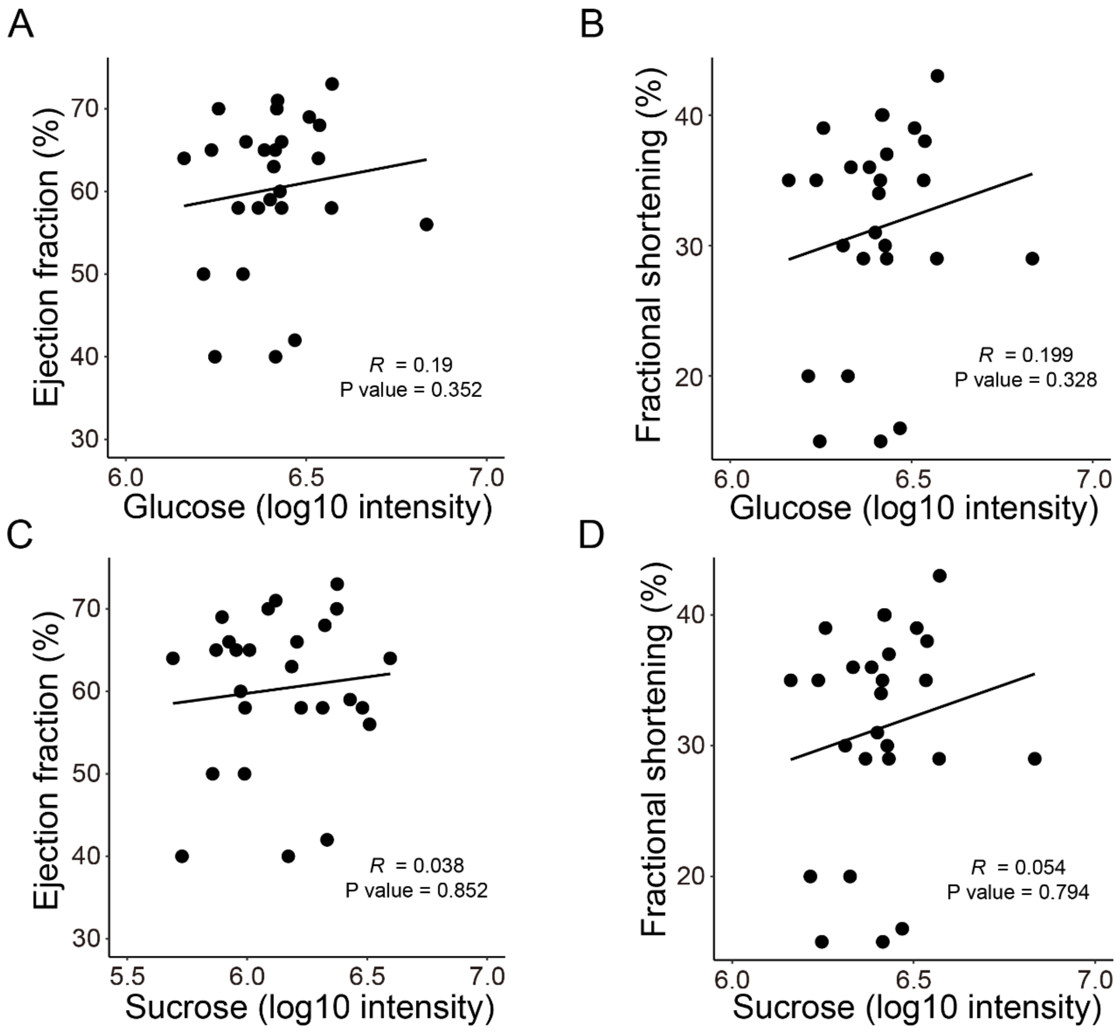


Figure S9. Glucose or sucrose levels were not associated with heart dysfunction in aortic stenosis patients. The ejection fraction and fractional shortening were not changed in aortic stenosis patients with a higher concentration of plasma (A) and (B) glucose or (C) and (D) sucrose, respectively (n=27).

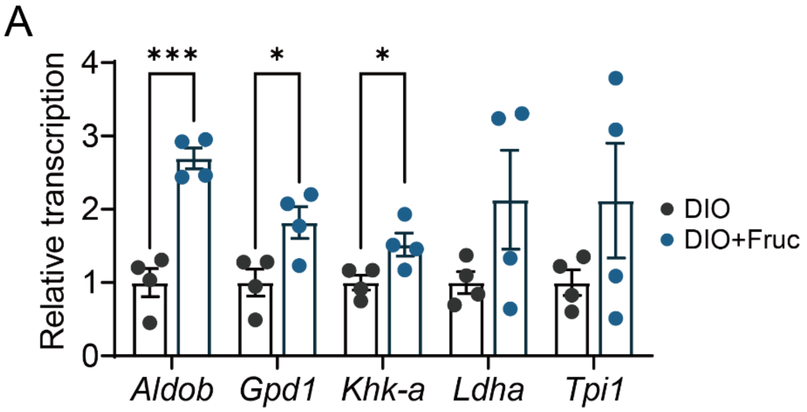


Figure S10. Fructolysis associated genes were upregulated in DIO+Fruc mouse (vs. DIO, *p<0.05, ***p<0.001, n=3).
