## Supplemental Tables for "Excessive Dietary Fructose Aggravates Heart Failure via Impairing Myocardial Fatty Acid Oxidation Metabolism in Diet Induced Obese Mouse"

1. Shanghai Key Laboratory of Metabolic Remodeling and Health, Institute of Metabolism and Integrative Biology and School of Life Sciences, Fudan University, Shanghai, 200438, China;
2. Department of Cardiology, Institute of Vascular Medicine, Peking University Third Hospital, NHC Key Laboratory of Cardiovascular Molecular Biology and Regulatory Peptides, State Key Laboratory of Vascular Homeostasis and Remodeling Peking University, Beijing, 100191, China;
3. School of Life Sciences, Inner Mongolia University, Hohhot Inner Mongolia, 010021, China;
4. Beijing Institute of Heart, Lung and Blood Vessel Diseases, Beijing Anzhen Hospital, Capital Medical University, Beijing 100029, China;
5. Department of Cardiology, Peking University First Hospital, Beijing, 100034, China;
6. Shanghai Institute of Nutrition and Health, Chinese Academy of Sciences, Shanghai, 200031, China;
7. Lead contact;

### These authors contribute equally

**Supplementary Table 1. The baseline characteristics of low (n=13) and high (n=14) fructose group.**

| Group | Low fructose (<0.021 mmol/L) | High fructose (≥0.021 mmol/L) | P value |
|---|---|---|---|
| Age, y | 68.62 (10.79) | 56.15 (8.99) | 0.004 |
| Female sex, n (%) | 3 (23.1) | 5 (38.5) | 0.671 |
| BMI, kg/$m^2$ | 25.05 (3.82) | 26.85 (4.17) | 0.261 |
| SBP, mmHg | 127.62 (22.65) | 115.23 (14.98) | 0.113 |
| Glucose, mmol/L | 5.65 (1.27) | 5.11 (0.81) | 0.170 |
| ALT, U/L | 16.31 (12.65) | 23.62 (20.78) | 0.290 |
| LDH, U/L | 199.75 (37.38) | 208.08 (40.37) | 0.599 |
| TC, mmol/L | 4.19 (1.00) | 5.02 (1.15) | 0.065 |
| LVEF, % | 64.77 (5.28) | 55.57 (10.15) | 0.007 |
| FS, % | 35.38 (4.54) | 26.93 (8.73) | 0.005 |
| eGFR, ml/min | 91.23 (14.62) | 80.48 (16.20) | 0.088 |
| Diuretics, n (%) | 9 (69.2) | 6 (46.2) | 0.427 |

BMI, body mass index; SBP, systolic blood pressure; ALT, alanine transaminase; LDH, lactate Dehydrogenase; TC, total cholesterol; LVEF, left ventricular ejection fraction; FS, fractional shortening; eGFR, estimated glomerular filtration rate.

**Supplementary Table 2: Association of BMI (Kg/m2) with incident various cardiovascular diseases.**

| | BMI | | |
|---|---|---|---|
| **Cardiovascular diseases** | **Odds ratio** | **95% CI** | **P value** |
| Hypertension | | | |
| Model 1 | 1.085 | [1.079, 1.090] | <0.0001 |
| Model 2 | 1.085 | [1.079, 1.091] | <0.0001 |
| Model 3 | 1.067 | [1.060, 1.075] | <0.0001 |
| Coronary artery disease | | | |
| Model 1 | 1.043 | [1.033, 1.054] | <0.0001 |
| Model 2 | 1.045 | [1.034, 1.056] | <0.0001 |
| Model 4 | 0.994 | [0.979, 1.008] | 0.387 |
| Heart failure | | | |
| Model 1 | 1.077 | [1.065, 1.090] | <0.0001 |
| Model 2 | 1.080 | [1.067, 1.092] | <0.0001 |
| Model 5 | 1.054 | [1.040, 1.068] | <0.0001 |

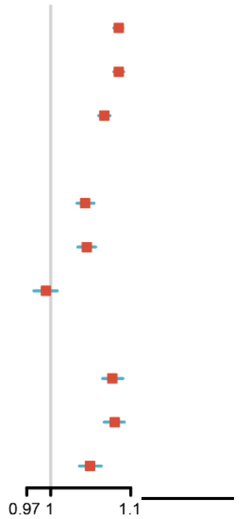

CI = confidence interval

Results were obtained from multinomial logistics regression models.

Model 1 adjusted for age, gender, and total energy intake.

Model 2 adjusted for model 1 plus race, education, marriage status, smoking status, total vegetable intake (cup), total grain intake (ounce), total dairy intake (cup), total protein intake (g), solid fat intake (g).

Model 3 adjusted for model 2 plus diabetes mellitus, heart failure, hyperlipidemia, coronary artery disease, blood glucose (mmol/l), blood total cholesterol (mmol/l), high density lipoprotein cholesterol (mml/l), lipid lowering treatment, anti-hypertensive drug, and anti-diabetic drug.

Model 4 adjusted for model 2 plus diabetes mellitus, hypertension, hyperlipidemia, heart failure, blood glucose (mmol/l), blood total cholesterol (mmol/l), high density lipoprotein cholesterol (mml/l), lipid lowering treatment, anti-hypertensive drug, and anti-diabetic drug.

Model 5 adjusted for model 2 plus diabetes mellitus, hypertension, hyperlipidemia, coronary artery disease, blood glucose (mmol/l), blood total cholesterol (mmol/l), high density lipoprotein cholesterol (mml/l), lipid lowering treatment, anti-hypertensive drug, and anti-diabetic drug.

**Supplementary Table 3: Association between percentage energy from macronutrients and cardiovascular disease.**

| | Odds ratio | 95% CI | P value |
|---|---|---|---|
| Percentage energy from carbohydrate (%) | | | |
| Hypertension | 0.999 | [0.995, 1.002] | 0.443 |
| Coronary artery disease | 1.009 | [1.001, 1.016] | 0.027 |
| Heart failure | 1.008 | [1.001, 1.016] | 0.035 |
| Percentage energy from total protein (%) | | | |
| Hypertension | 0.992 | [0.985, 1.000] | 0.053 |
| Coronary artery disease | 0.987 | [0.974, 1.001] | 0.075 |
| Heart failure | 0.991 | [0.975, 1.008] | 0.301 |
| Percentage energy from total fat (%) | | | |
| Hypertension | 0.992 | [0.988, 0.996] | <0.001 |
| Coronary artery disease | 0.993 | [0.984, 1.002] | 0.104 |
| Heart failure | 0.994 | [0.986, 1.003] | 0.206 |

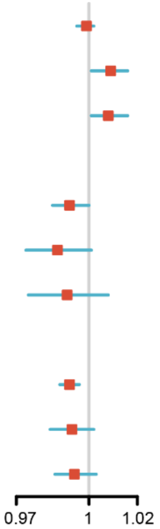


CI = confidence interval

Results were obtained from multinomial logistics regression models, including age, sex, race, education, marriage status, body mass index (Kg/$m^2$), smoking status, diabetes, antihypertensive medication use, Healthy Eating Index score, total serum cholesterol, serum glucose, and total energy intake.

**Supplementary Table 4: Association of total sugar intake (g) per day with incident various cardiovascular diseases.**

| | Total sugar intake | | |
|---|---|---|---|
| **Cardiovascular diseases** | **Odds ratio** | **95% CI** | **P value** |
| Hypertension | | | |
| Model 1 | 0.999 | [0.999, 1.000] | 0.055 |
| Model 2 | 0.999 | [0.998, 0.999] | <0.001 |
| Model 3 | 0.999 | [0.998, 1.000] | 0.001 |
| Coronary artery disease | | | |
| Model 1 | 1.000 | [0.998, 1.001 | 0.718 |
| Model 2 | 1.001 | [0.999, 1.003] | 0.263 |
| Model 4 | 1.001 | [0.999, 1.003] | 0.246 |
| Heart failure | | | |
| Model 1 | 1.002 | [1.000, 1.003] | 0.053 |
| Model 2 | 1.002 | [1.000, 1.004] | 0.024 |
| Model 5 | 1.002 | [1.000, 1.004] | 0.023 |

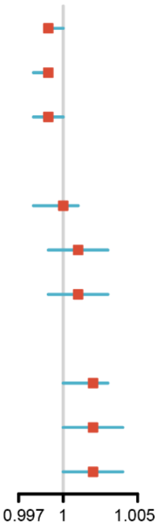


CI = confidence interval

Results were obtained from multinomial logistics regression models.

Model 1 adjusted for age, gender, body mass index (Kg/m$^2$), and total energy intake.

Model 2 adjusted for model 1 plus race, education, marriage status, smoking status, total vegetable intake (cup), total grain intake (ounce), total dairy intake (cup), total protein intake (g), solid fat intake (g).

Model 3 adjusted for model 2 plus diabetes mellitus, heart failure, hyperlipidemia, coronary artery disease, blood glucose (mmol/l), blood total cholesterol (mmol/l), high density lipoprotein cholesterol (mml/l), lipid lowering treatment, anti-hypertensive drug, and anti-diabetic drug.

Model 4 adjusted for model 2 plus diabetes mellitus, hypertension, hyperlipidemia, heart failure, blood glucose (mmol/l), blood total cholesterol (mmol/l), high density lipoprotein cholesterol (mml/l), lipid lowering treatment, anti-hypertensive drug, and anti-diabetic drug.

Model 5 adjusted for model 2 plus diabetes mellitus, hypertension, hyperlipidemia, coronary artery disease, blood glucose (mmol/l), blood total cholesterol (mmol/l), high density lipoprotein cholesterol (mml/l), lipid lowering treatment, anti-hypertensive drug, and anti-diabetic drug.

**Supplementary Table 5. Sensitivity analysis for the causal association between blood fructose and cardiovascular disease.**

| **Cardiovascular diseases** | **Method** | **No. of SNPs** | **OR (95% CI)** | **P value of association** | **P-intercept** |
|---|---|---|---|---|---|
| Hypertension | | | | | |
| | MR-PRESSO test | 29 | 0.97 (0.76, 1.24) | 0.81 | 0.89 |
| | Maximum likelihood based | 29 | 0.97 (0.75, 1.25) | 0.81 | |

| | | | | | |
|---|---|---|---|---|---|
| | MR-Egger regression | 29 | 1.00 (0.58, 1.75) | 0.97 | |
| | Weighted median | 29 | 0.93 (0.67, 1.29) | 0.67 | |
| Coronary artery disease | | | | | |
| | MR-PRESSO test | 24 | 1.11 (0.97, 1.25) | 0.14 | 0.057 |
| | Maximum likelihood based | 24 | 1.10 (0.92, 1.32) | 0.28 | |
| | MR-Egger regression | 24 | 1.56 (1.05, 2.32) | 0.027 | |
| | Weighted median | 24 | 1.20 (1.01, 1.43) | 0.036 | |
| Heart failure | | | | | |
| | MR-PRESSO test | 25 | 1.21 (1.04, 1.42) | 0.021 | 0.56 |
| | Maximum likelihood based | 25 | 1.49 (1.10, 2.02) | 0.0073 | |
| | MR-Egger regression | 25 | 1.57 (0.82, 3.02) | 0.17 | |
| | Weighted median | 25 | 1.10 (0.89, 1.35) | 0.37 | |

OR = odds ratio, CI = confidence interval

**Supplementary Table 6. Sensitivity analysis for the causal association between blood glucose and cardiovascular disease.**

| Cardiovascular diseases | Method | No. of SNPs | OR (95% CI) | P value of association | P-intercept |
|---|---|---|---|---|---|
| Hypertension | | | | | |
| | MR-PRESSO test | 111 | 0.95 (0.67, 1.35) | 0.79 | 0.23 |
| | Maximum likelihood based | 111 | 0.95 (0.66, 1.36) | 0.78 | |
| | MR-Egger regression | 111 | 0.69 (0.36, 1.30) | 0.25 | |
| | Weighted median | 111 | 0.84 (0.50, 1.42) | 0.52 | |
| Coronary artery disease | | | | | |
| | MR-PRESSO test | 101 | 1.13 (0.93, 1.39) | 0.21 | 0.14 |
| | Maximum likelihood based | 101 | 1.14 (0.93, 1.41) | 0.21 | |
| | MR-Egger regression | 101 | 0.88 (0.59, 1.30) | 0.52 | |
| | Weighted median | 101 | 1.08 (0.82, 1.42) | 0.57 | |
| Heart failure | | | | | |
| | MR-PRESSO test | 101 | 1.01 (0.80, 1.26) | 0.81 | 0.84 |
| | Maximum likelihood based | 101 | 1.01 (0.80, 1.27) | 0.94 | |
| | MR-Egger regression | 101 | 1.05 (0.65, 1.69) | 0.83 | |
| | Weighted median | 101 | 0.97 (0.72, 1.33) | 0.87 | |

OR = odds ratio, CI = confidence interval

**Supplementary Table 7. Sensitivity analysis for the causal association between blood sucrose and cardiovascular disease.**

| Cardiovascular diseases | Method | No. of SNPs | OR (95% CI) | P value of association | P-intercept |
|---|---|---|---|---|---|
| Hypertension | | | | | |
| | MR-PRESSO test | 32 | 1.00 (0.99, 1.01) | 0.93 | 0.088 |
| | Maximum likelihood based | 32 | 0.95 (0.66, 1.36) | 0.78 | |
| | MR-Egger regression | 32 | 1.00 (0.99, 1.01) | 0.93 | |
| | Weighted median | 32 | 0.99 (0.98, 1.01) | 0.85 | |
| Coronary artery disease | | | | | |
| | MR-PRESSO test | 26 | 1.00 (1.00, 1.01) | 0.55 | 0.57 |
| | Maximum likelihood based | 26 | 1.00 (0.99, 1.00) | 0.60 | |
| | MR-Egger regression | 26 | 1.00 (0.99, 1.01) | 0.76 | |
| | Weighted median | 26 | 0.999 (0.995, 1.003) | 0.91 | |
| Heart failure | | | | | |
| | MR-PRESSO test | 24 | 1.00 (0.99, 1.00) | 0.056 | 0.086 |
| | Maximum likelihood based | 24 | 0.99 (0.99, 1.00) | 0.073 | |
| | MR-Egger regression | 24 | 0.990 (0.982, 0.999) | 0.022 | |
| | Weighted median | 24 | 0.995 (0.991, 1.000) | 0.054 | |

OR = odds ratio, CI = confidence interval

**Supplementary Table 8. Bidirectional analysis for the causal association of heart failure with fructose/glucose.**

| Heart failure | Method | No. of SNPs | OR (95% CI) | P value of association | P-intercept |
|---|---|---|---|---|---|
| Fructose | | | | | |
| | MR-PRESSO test | 130 | 1.00 (0.98, 1.02) | 0.86 | |
| | Maximum likelihood based | 130 | 1.00 (0.98, 1.02) | 0.85 | |
| | MR-Egger regression | 130 | 1.03 (0.95, 1.11) | 0.47 | 0.56 |
| | Weighted median | 130 | 1.00 (0.97, 1.02) | 0.82 | |
| | Inverse variance weighted | 130 | 1.00 (0.98, 1.02) | 0.86 | |
| Glucose | | | | | |
| | MR-PRESSO test | 130 | 1.01 (1.00, 1.02) | 0.0035 | |
| | Maximum likelihood based | 130 | 1.01 (1.00, 1.02) | 0.0052 | |
| | MR-Egger regression | 130 | 1.01 (0.98, 1.04) | 0.41 | 0.57 |
| | Weighted median | 130 | 1.01 (0.99, 1.02) | 0.17 | |
| | Inverse variance weighted | 130 | 1.01 (1.00. 1.02) | 0.0049 | |

OR = odds ratio, CI = confidence interval

**Supplementary Table 9. Results for the Mendelian randomization heterogeneity analysis under Inverse variance weighted method.**

| Exposure | Outcome | Heterogeneity Q value ($I^2$) | P value |
|---|---|---|---|
| Fructose | Hypertension | 36.19 (22.64%) | 0.14 |
| | Coronary artery disease | 26.98 (14.75%) | 0.26 |
| | Heart failure | 29.66 (19.08%) | 0.2 |
| Glucose | Hypertension | 131.61 (16.42%) | 0.079 |
| | Coronary artery disease | 136.23 (26.59%) | 0.0094 |
| | Heart failure | 136.23 (26.59%) | 0.0094 |
| Heart failure | Fructose | 115.34 (11.85%) | 0.8 |
| | Glucose | 1.01 (1.00, 1.02) | 0.0035 |
